# The Mettl3 epitranscriptomic writer amplifies p53 stress responses

**DOI:** 10.1101/2021.10.21.465324

**Authors:** Nitin Raj, Mengxiong Wang, Jose A. Seoane, Nancie A. Moonie, Janos Demeter, Alyssa M. Kaiser, Anthony M. Boutelle, Abigail S. Mulligan, Clare Moffatt, Christina Curtis, Howard Y. Chang, Peter. K. Jackson, Laura D. Attardi

## Abstract

The p53 transcription factor, encoded by the most frequently mutated gene in human cancer, plays a critical role in tissue homeostasis in response to stress signals. The mechanisms through which p53 promotes downstream tumor suppressive gene expression programs remain, however, only superficially understood. Here, we used tandem affinity purification and mass spectrometry to reveal new components of the p53 response. This approach uncovered Mettl3, a component of the m^6^A RNA methyltransferase complex (MTC), as a p53-interacting protein. Analysis of Mettl3- deficient cells revealed that Mettl3 promotes p53 protein stabilization and target gene expression in response to DNA damage. Mettl3 acts in part by competing with the p53 negative regulator, Mdm2, for binding to the p53 transactivation domains to promote methyltransferase-independent stabilization of p53. In addition, Mettl3 relies on its catalytic activity to augment p53 responses, with p53 recruiting Mettl3 to p53 target genes to co-transcriptionally direct m^6^A modification of p53 pathway transcripts to enhance their expression. Mettl3 also promotes p53 activity downstream of oncogenic signals *in vivo*, in both allograft and autochthonous lung adenocarcinoma models, suggesting cooperative action of p53 and Mettl3 in tumor suppression. Accordingly, we found in diverse human cancers that mutations in MTC components perturb expression of p53 target genes and that MTC mutations are mutually exclusive with *TP53* mutations, suggesting that the MTC enhances the p53 transcriptional program in human cancer. Together, these studies reveal a fundamental role for Mettl3 in amplifying p53 signaling through protein stabilization and epitranscriptome regulation.

## INTRODUCTION

The *TP53* gene, which encodes the p53 protein, is mutated in over half of all human cancers, reflecting its fundamental role as a tumor suppressor^1^. p53 is a transcription factor that integrates cellular stress cues such as DNA damage and oncogenic signaling to drive anti-proliferative or pro-apoptotic responses important for tissue homeostasis and tumor suppression^2–4^. p53 activity as a transcriptional activator is critical for governing gene expression programs that limit tumorigenesis, as evidenced by the preponderance of *TP53* mutations observed in sequences encoding the DNA binding domain in human cancers and studies of p53 transactivation dead mutant knock-in mice, which are highly tumor prone^5, 6^. Despite this well-established role for p53 transcriptional function in tissue homeostasis in response to cellular stresses, however, the mechanisms through which p53 regulates a downstream expression program remain only superficially understood. A complete understanding of the mechanisms of p53-regulated gene expression are of paramount importance for ultimately targeting this critical tumor suppressor in the clinic^7^.

p53 promotes specific gene expression programs through recruitment of the general transcriptional machinery via cofactors such as mediator subunits and TAFs to increase levels of transcriptional initiation^8, 9^. p53 also recruits histone-modifying enzymes, including p300, CBP, and PCAF, to acetylate histones and remodel chromatin^10^. However, the mechanisms by which p53 regulates gene expression beyond effects on transcriptional initiation remain only superficially understood. Deciphering additional aspects of p53 activity will provide key insight into p53 function in tissue homeostasis and cancer suppression.

Eukaryotic gene expression is regulated at multiple levels to provide maximal capacity for fine-tuning in response to various cues such as DNA damage and oncogenic signaling^11^. The ultimate gene expression profile of a cell can be influenced by modulation of transcription, RNA metabolism, and protein translation or stability. While RNA metabolism has been studied at the level of splicing for many years, recent work has illuminated the importance of RNA modification in regulation of gene expression. Specifically, N(6)-methyladenosine (m^6^A), the most abundant and prevalent internal modification in eukaryotic mRNAs, is installed on mRNAs co-transcriptionally, primarily by the Mettl3-Mettl14 methyltransferase writer complex. Depending on the context and the m^6^A readers expressed in a given setting, m^6^A modification can then affect gene expression through effects on RNA stability, subcellular localization, and translation. Importantly, m^6^A modification has been demonstrated to be critical for cell state transitions, such as during differentiation of stem cells and responses to a variety of stress signals, such as DNA damage or heat shock^12^. However, the factors that govern which transcripts are selected for m^6^A modification to drive specific cellular responses remain enigmatic.

Here, we seek to gain new insight into how p53 regulates gene expression programs in the context of stress signals using tandem affinity purification coupled with mass spectrometry to identify novel p53-interacting partners. To ensure that p53 pathways remain intact, we perform experiments using untransformed cells, and we identify the Mettl3 m^6^A RNA-methyltransferase as a p53-interacting protein. Remarkably, analysis of Mettl3-deficient cells reveals that both p53 stabilization and p53 target gene induction by either DNA damage or oncogenic signals are significantly compromised by Mettl3 deficiency. Interestingly, Mettl3 binds to the same region of p53 as the Mdm2 ubiquitin ligase, and therefore effectively competes with Mdm2, thus providing key insight into how p53 stability is enhanced by Mettl3. Moreover, Mettl3 binding to p53 not only stabilizes p53 but also co-transcriptionally directs m^6^A modification of downstream components of the p53 pathway. We find further that Mettl3 not only reinforces p53 function in response to DNA damage but also in tumor suppression, both in mice and humans. Collectively, these studies reveal a fundamental role for Mettl3 in amplifying p53 signaling both through protein stabilization and epitranscriptome regulation.

## RESULTS

### Mettl3 interacts with p53 and enhances p53 activity

To identify novel p53-interacting partners, we expressed LAP (Localization and affinity purification)-tagged p53 in NIH-3T3 fibroblasts and treated cells with the DNA double strand break-inducer doxorubicin (dox), which triggers p53-dependent G1 arrest, providing a good model for understanding the biochemical basis for p53 action^13^ (data not shown). After purification and mass spectrometry, we identified Mettl3, the catalytic component of the core methyltransferase complex that installs *N6*-methyladenosine (m^6^A) on eukaryotic messenger RNAs (mRNAs), as a p53-interacting partner in both basal and DNA damage conditions (Fig.1a, b, Raj et al., in preparation)^12^. We confirmed the p53-Mettl3 interaction by additional co-immunoprecipitation assays with tagged variants of the proteins as well as with endogenous proteins in primary mouse embryonic fibroblasts (MEFs; Fig. 1c, Extended Data Fig. 1a). Through co-immunoprecipitation assays in fibroblasts, we found further that p53 can also interact with Mettl14, a heterodimeric partner of Mettl3, suggesting that p53 interacts with the methyltransferase complex critical for m^6^A RNA modification (Extended Data Fig. 1b)^14, 15^. The Mettl3-p53 interaction is not indirectly mediated by DNA, as the interaction is still observed upon ethidium bromide or DNAse treatment (Extended Data Fig. 1c, d).

**Fig. 1.**
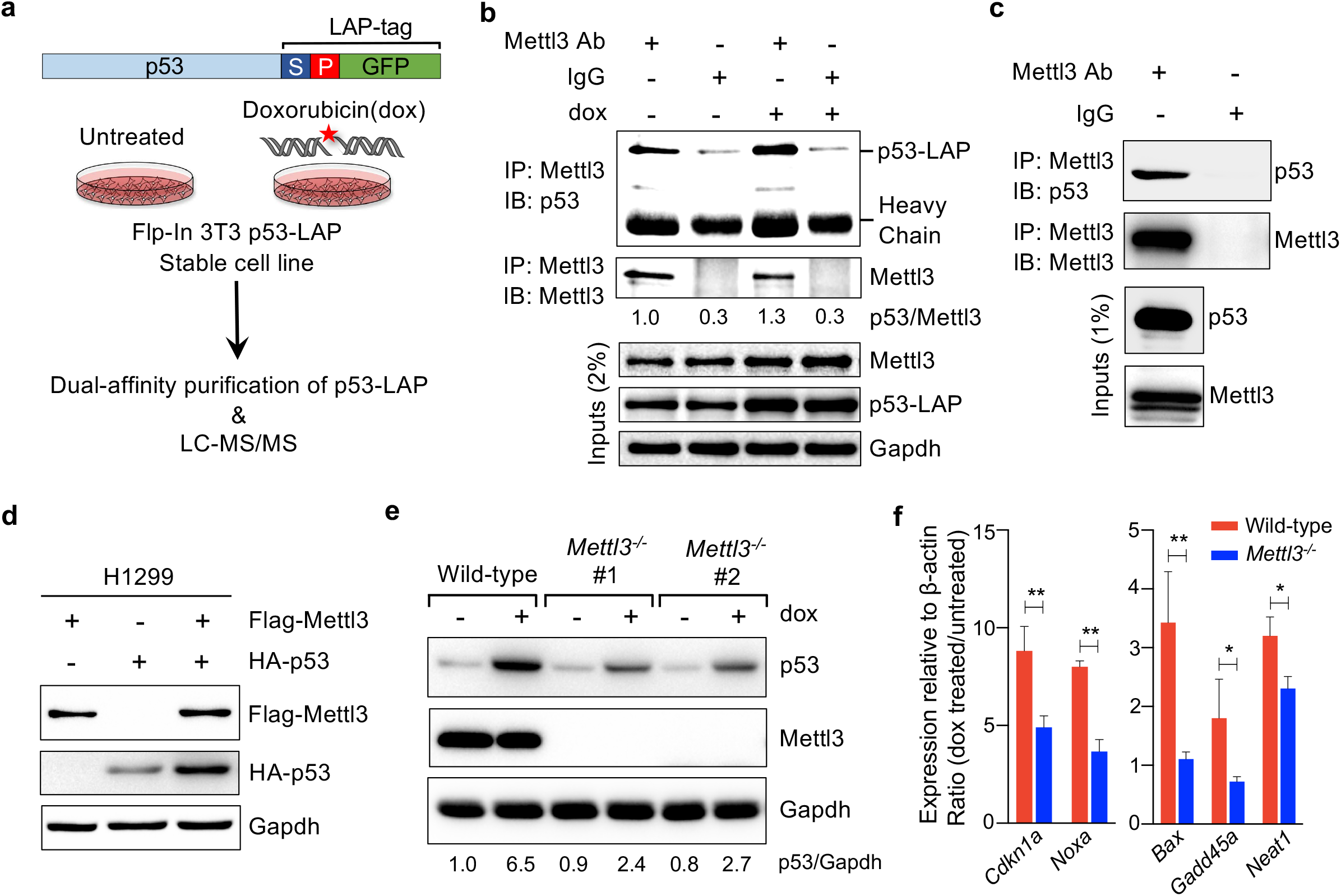
Mettl3 interacts with p53 and enhances p53 transcriptional activity in DNA- damage treated cells. **a,** Schematic of dual-affinity purification of LAP-tagged p53 protein in Flp-In-3T3 fibroblasts. Cells were either left untreated or treated with 0.2 μg/mL doxorubicin (dox) for 6 hours, followed by dual affinity purification of p53-bound protein complexes and protein identification by LC-MS/MS. **b,** Co-immunoprecipitation (Co-IP) and immunoblot assay to validate p53-LAP and endogenous Mettl3 interaction in untreated and DNA damage treated Flp-In-3T3 cells. Numbers underneath indicate the amount of p53 co-precipitated relative to Mettl3 (*n*=3). **c,** Co-IP and immunoblot assay to examine interaction of endogenous Mettl3 and p53 in *E1A;HRasV12*-expressing MEFs (*n*=2). **d,** Immunoblot after transfection of Flag-Mettl3 and HA-p53 plasmids into H1299 cells (*n*=3). **e,** p53 induction after 6 hours dox (0.2 μg/mL) in wild-type and two different *Mettl3^-/-^* ES cell lines. Numbers underneath indicate the amount of p53 relative to Gapdh loading control (*n*=3). **f**, Induction of p53 target genes after 6 hours dox (0.2 μg/mL) in *Mettl3^-/-^* ES cells relative to wild-type cells, normalized to *β-Actin*. *Mettl3^-/-^* depicts mean of two different *Mettl3* knockout cell lines. Data are mean ± s.e.m. of at least three biological replicates each with three technical replicates. *P* values were determined by the unpaired two-tailed Student’s *t*-test. \**P*<0.05, \*\**P*<0.01. Representative immunoblots are shown in **b,c,d** and **e**, and Gapdh serves as a loading control.

Interestingly, upon co-expression of p53 and Mettl3 in p53-deficient human or mouse cells, we noted that Mettl3 enhances p53 protein levels, suggesting that Mettl3 might affect p53 function (Fig. 1d, Extended Data Fig. 2a). To test this hypothesis, we sought to assess the consequences of Mettl3 deficiency for p53 function. We first leveraged *Mettl3* null embryonic stem (ES) cells and assessed how Mettl3 loss affects p53 responses to DNA damage^16^. We found that *Mettl3* nullizygosity results in decreased p53 accumulation in response to DNA damage, which is accompanied by diminished induction of p53 target genes, indicating that Mettl3 enhances p53 protein accumulation and transcriptional activity in ES cells (Fig. 1e, f, Extended Data Fig. 2b). To determine whether this response extends to other cell types, we next examined oncogene-expressing primary MEFs. Similarly, we noted that p53 accumulation and target gene induction in response to DNA damage are impaired upon Mettl3 knockdown (Fig. 2a, b). Collectively, these findings indicate that Mettl3 enforces robust p53 responses to DNA damage.

**Fig. 2.**
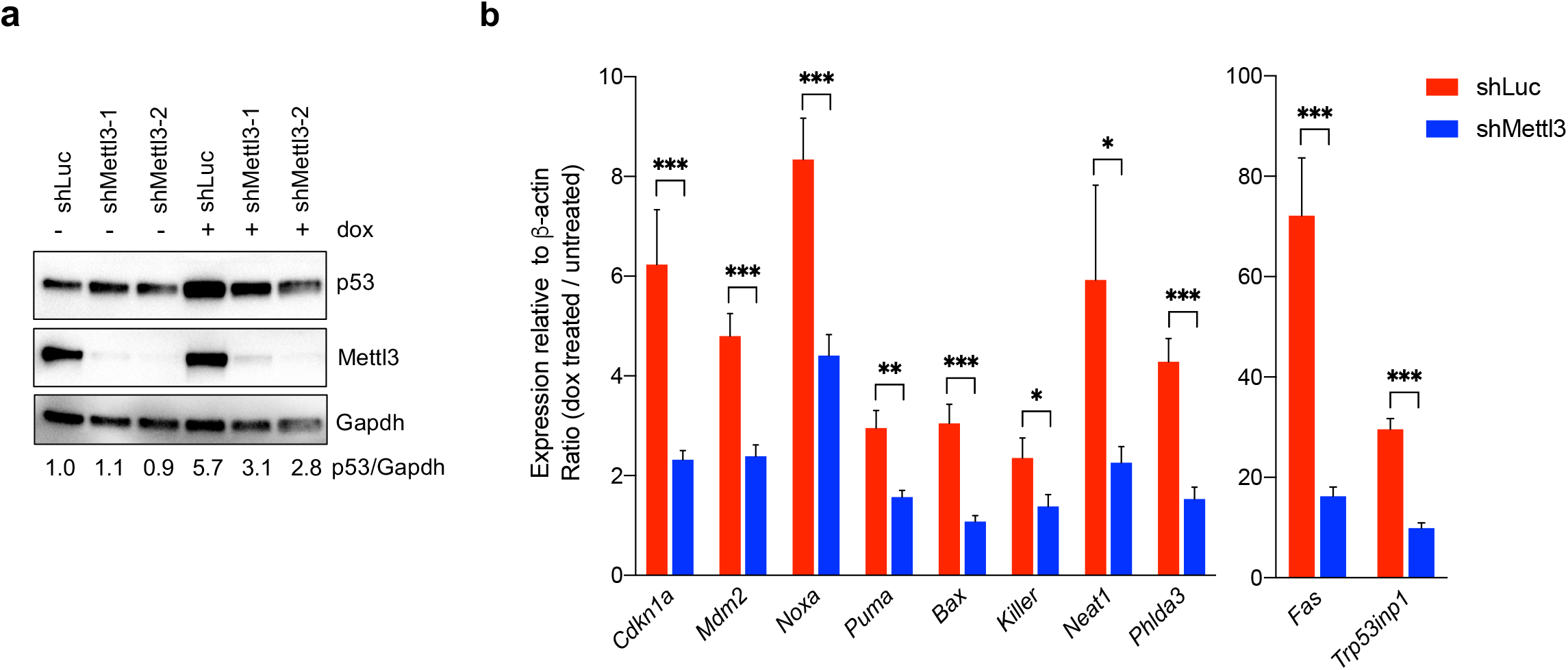
Mettl3 knockdown impairs p53 accumulation and target gene induction in response to DNA damage in oncogene-expressing primary MEFs. **a,** Immunoblots of *E1A;HRasV12*-expressing MEFs showing protein levels of p53 and Mettl3 in shLuc and shMettl3 shRNA-expressing cell lines. Gapdh serves as loading control. Representative immunoblots are shown from three biological replicates, and two different MEF lines per genotype were used for this experiment. The numbers indicate the p53 levels after normalization to Gapdh. **b,** qRT-PCR analysis of expression of p53 target genes after 6 hours dox (0.2 μg/mL) in shLuc and shMettl3 shRNA-expressing cell lines, normalized to *β-Actin*. shMettl3 depicts mean of cell lines expressing either of two different shRNAs targeting Mettl3. Representative data are shown from two biological replicates each with three technical replicates and two different MEF lines per genotype were used. Data are mean of fold induction (Ratio of dox treated / untreated) ± s.e.m. *P* values were determined by the unpaired two-tailed Student’s *t*-test. \**P*<0.05, \*\**P*<0.01, \*\*\**P*<0.001.

### Mettl3 promotes p53 stability through competition with Mdm2

We next investigated the molecular mechanisms by which Mettl3 enhances p53 protein levels. p53 protein has a short half-life resulting from interaction with the Mdm2 ubiquitin ligase, which targets p53 for ubiquitin-mediated proteolysis^17, 18^. In response to stress signals, Mdm2 is displaced from p53, resulting in enhanced p53 protein stability and activation of p53 transcriptional programs^19, 20^. We therefore sought first to assess whether Mettl3 might affect p53 protein stability. We treated ES cells with dox to stabilize p53, then with cycloheximide to block translation, and interrogated how Mettl3 status affects p53 half-life. Interestingly, Mettl3 deficiency was characterized by a diminished p53 half-life, suggesting that Mettl3 acts to enhance p53 protein stabilization (Fig. 3a, Extended Data Fig. 3). To assess whether the enhanced p53 stabilization by Mettl3 reflects differences in Mdm2 binding, we performed Co-IP experiments to measure the Mdm2-p53 interaction in the presence and absence of Mettl3. Intriguingly, in the absence of Mettl3, Mdm2 interacts more strongly with p53 upon DNA damage than when Mettl3 is expressed (Fig. 3b). These findings suggest that in response to DNA damage, Mdm2 is efficiently displaced from p53 when Mettl3 is present but not when it is absent.

**Fig. 3.**
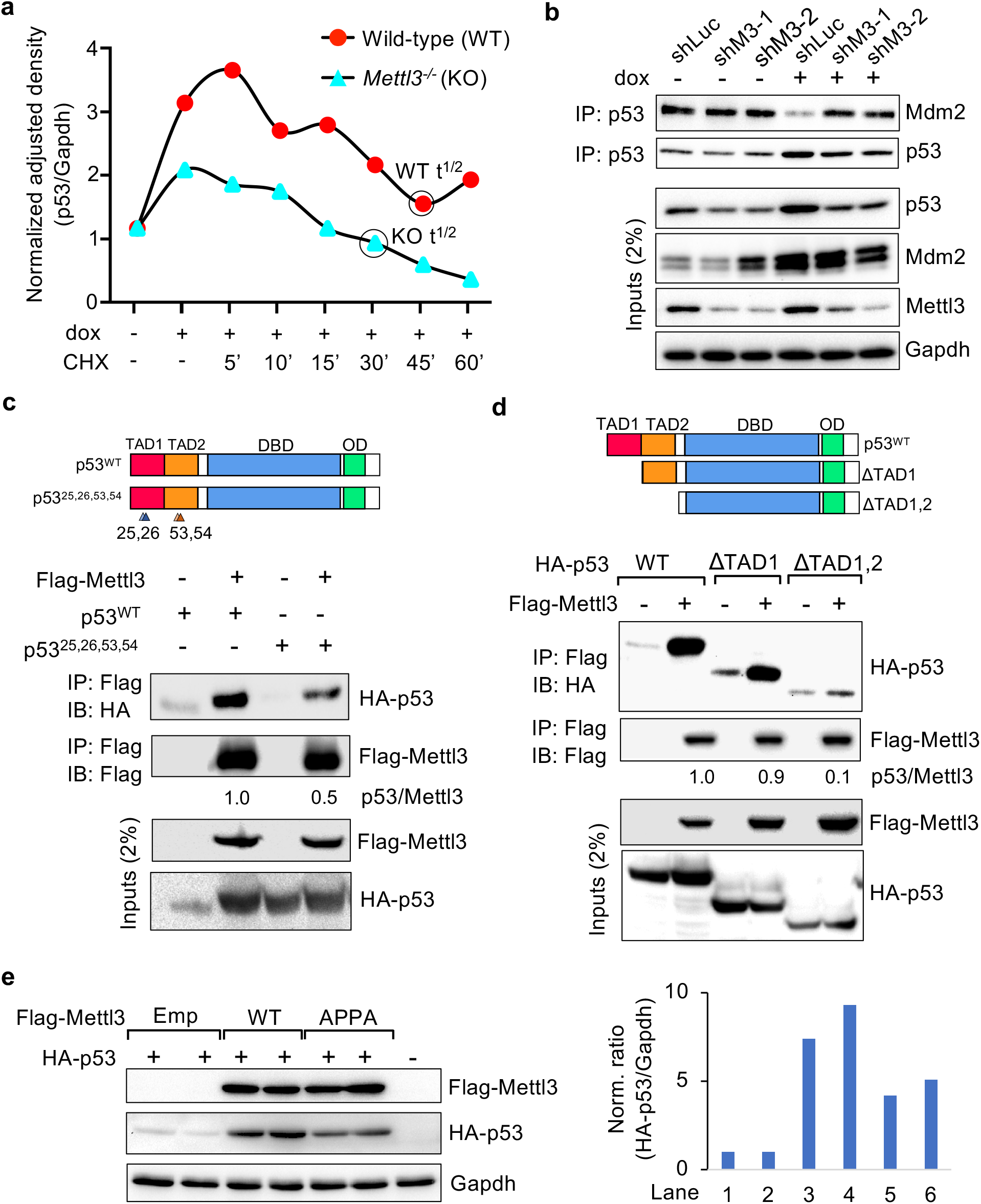
Mettl3 enhances p53 protein half-life by displacing Mdm2. **a,** Time course analysis of p53 levels, normalized to Gapdh, in wild-type (WT) and *Mettl3^-/-^* (KO) mouse ES cells after treatment with 0.2 μg/mL doxorubicin (dox) and cycloheximide (CHX) (*n*=2). **b,** Co-immunoprecipitation and immunoblot assays in *E1A;HRasV12*-expressing MEFs to assess Mdm2 pull-down with p53, in *E1A;HRasV12*-expressing MEFs expressing control shLuc or shMettl3 RNAs, under basal and DNA-damage conditions [6 hours dox (0.2 μg/mL)]. Representative immunoblots are shown from four biological replicates and two different MEF lines per genotype were used. Gapdh serves as a loading control. **c,** Co-immunoprecipitation and immunoblot assays in Flp-In-3T3 *p53^-/-^* cells to assess pull- down of HA-p53 wild-type or transcriptionally-dead HA-p53^25, 26, 53, 54^ (schematized at top) with Flag-Mettl3. Numbers indicate the amount of HA-p53 co-immunoprecipitated relative to Flag-Mettl3 (*n*=3). **d,** Co-immunoprecipitation and immunoblot assays in Flp-In-3T3 *p53^-/-^* cells to assess pull-down of HA-p53 wild-type or HA-p53 deletion mutants schematized at top with Flag-Mettl3. Numbers indicate the amount of HA-p53 co- immunoprecipitated relative to Flag-Mettl3 (*n*=3). **e,** (Left) Immunoblot analysis to assess p53 levels upon co-transfection of HA-p53 and empty vector, Flag-Mettl3 (WT) or catalytically inactive Flag-Mettl3 (APPA) into *p53^-/-^* Flp-In-3T3 cells. Gapdh serves as a loading control. (Right) Quantitation of HA-p53 levels relative to Gapdh. Representative immunoblots are shown in **c,d,** and **e**.

This observation suggested the possibility that Mettl3 and Mdm2 might compete for binding on p53. We therefore tested whether Mettl3 binds to the same region of p53 as Mdm2. Mdm2 binds to the p53 amino-terminus, where the two p53 transcriptional activation domains (TADs) reside between residues 1-80. The interaction of Mdm2 with p53 is in fact disrupted by mutations in the two TADs at residues L25,Q26 in TAD1 and F53,F54 in TAD2^10, 21^. Notably, the ability of Mettl3 to bind to p53 is reduced by ∼50% in the p53^L25Q,Q26S,F53Q,F54S^ mutant (Fig. 3c). Moreover, deletion of both p53 TADs virtually abolished the interaction with Mettl3 (Fig. 3d). These observations suggest that Mettl3 binds to the p53 TADs and support the idea that Mettl3 stabilizes p53 by hindering Mdm2 binding. If so, then the effects of Mettl3 on p53 might be expected to be at least partially independent of its catalytic activity. To test this hypothesis, we used a Mettl3 catalytic mutant (Mettl3^D395A,W398A^) that lacks m^6^A methyltransferase activity^22^. Indeed, co-expression of this catalytically-dead Mettl3 mutant with p53 results in efficient enhancement of p53 stabilization, albeit somewhat less than with wild-type Mettl3 (Fig. 3e). Together, these findings reveal that Mettl3 plays a catalytic activity-independent role to stabilize p53 but also suggest that Mettl3 might exert a catalytic activity-dependent role in augmenting the p53 pathway.

### Mettl3 promotes m^6^A modification of p53 pathway transcripts

To interrogate the catalytic activity-dependent role of Mettl3 in the p53 pathway, we performed m^6^A-eCLIP-seq to identify those Mettl3-dependent m^6^A RNA modifications occurring after a stress signal. We utilized oncogene-expressing fibroblasts expressing control shRNAs or shRNAs directed against Mettl3 to acutely knockdown Mettl3, and we either left cells untreated or treated 6 hours with dox before performing m^6^A-eCLIP-seq (Fig. 4a, b). We first queried the site of m^6^A modifications in each condition and found that m^6^A modifications mapped primarily to the classical DRACH motif and were predominantly localized to the 3’ UTR or last exon of the coding sequence of mRNAs (Fig. 4c, d, Extended Data Fig. 4a, b). We next characterized the transcripts displaying Mettl3-dependent m^6^A modification in the presence of DNA damage when p53 function is maximal. After careful normalization of m^6^A IP signal to input to ensure analysis of differences in m^6^A modification between samples, functional annotation analysis of the transcripts with enhanced Mettl3-dependent m^6^A modification under DNA damaging agent conditions revealed p53-bound and p53-regulated transcripts as among the top identified categories (Fig. 4e). The overlap between the genes that undergo Mettl3- dependent m^6^A modification under DNA damage conditions and genes found previously to be bound and regulated by p53 upon DNA damage in MEFs^23^ was statistically significant (Fisher exact test, *P*=5.9e-016). The numerous p53 target gene transcripts with Mettl3-dependent m^6^A modification upon DNA damaging agent treatment included *Trp53inp1, Mdm2, Ccng1* and *Pmaip (Noxa)* (Fig 4f, g). In contrast, no clear Mettl3- dependent m^6^A modification was detected for the *p53* transcript itself upon treating with DNA damage (data not shown).

**Fig. 4.**
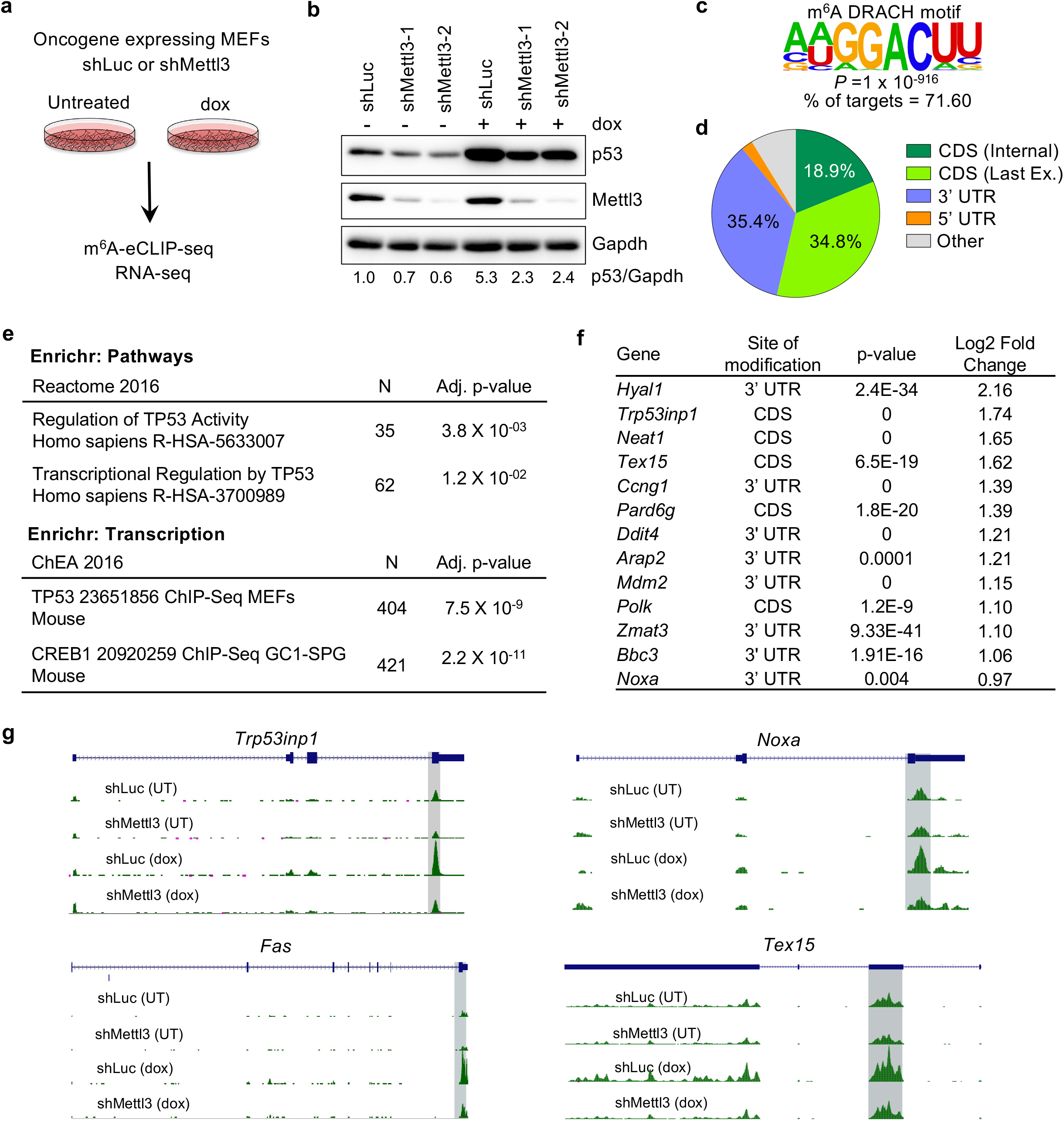
Mettl3-mediated m^6^A modification regulates p53 signaling pathway. **a,** *E1A;HRasV12*-expressing wild-type MEFs transduced with shLuc or shMettl3 hairpins were either left untreated or treated with 0.2 μg/mL doxorubicin (dox) for 6 hours followed by m^6^A-eCLIP-seq and RNA-seq analysis. **b**, Immunoblots showing p53 and Mettl3 protein levels in shLuc and shMettl3 shRNA-expressing cell lines. Gapdh serves as a loading control. Representative immunoblots are shown from three biological replicates, and two different MEF lines per genotype were used for this experiment. **c,** Identification of the known consensus m^6^A DRACH motif in mRNAs displaying m^6^A modification in the presence of DNA damage, by performing *de novo* motif search with HOMER database. **d,** Pie chart of the distribution of m^6^A peaks enriched upon DNA damage. m^6^A-IP reads were normalized to the total number of reads covering the m^6^A residue in the input sample. **e,** Functional annotation analysis of Mettl3-dependent m^6^A peaks enriched upon DNA-damage using Enrichr (N=number of genes per term). **f,** Table of p53 pathway transcripts with Mettl3-dependent m^6^A modification under acute DNA damage showing major site of modification, *p*-values and log2 fold change for enriched m^6^A peaks. **g,** UCSC genome browser tracks showing RPM (reads per million) patterns of m^6^A-eCLIP-seq in *Trp53inp1, Noxa, Fas and Tex15* mRNAs in untreated or doxorubicin-treated *E1A;HRasV12* MEFs transduced with either shLuc or shMettl3 RNAs.

The Mettl3-dependent modification of myriad p53 target gene transcripts upon DNA damage suggests that the p53-Mettl3 interaction augments the p53-driven gene expression program in part through co-transcriptional installation of m^6^A on transcripts. To elucidate the mechanism underlying this effect of Mettl3, we examined whether Mettl3 might associate with chromatin at p53 target genes. Using ChIP assays in MEFs, we found that Mettl3 associates with the p53 binding sites of p53 target genes, including *Mdm2* and *Noxa*, suggesting that Mettl3 can associate with chromatin of p53 target genes to direct m^6^A modification (Fig. 5a). This interaction is diminished in the absence of p53, underscoring the importance of p53 for Mettl3 recruitment to these sites (Fig. 5b). Together, these findings suggest that the Mettl3 complex installs m^6^A marks on select p53-regulated mRNAs co-transcriptionally. Analysis of the *Noxa* 3’UTR fused to a heterologous luciferase reporter revealed that Mettl3 promotes reporter expression, suggesting that m^6^A modification has the potential to enhance expression of p53 pathway transcripts (Fig. 5c, d). Overall, our findings suggest that Mettl3 exerts a dual effect on p53, through both stabilization of p53 and through modification of various transcripts in the p53 pathway to enhance their expression.

**Fig. 5.**
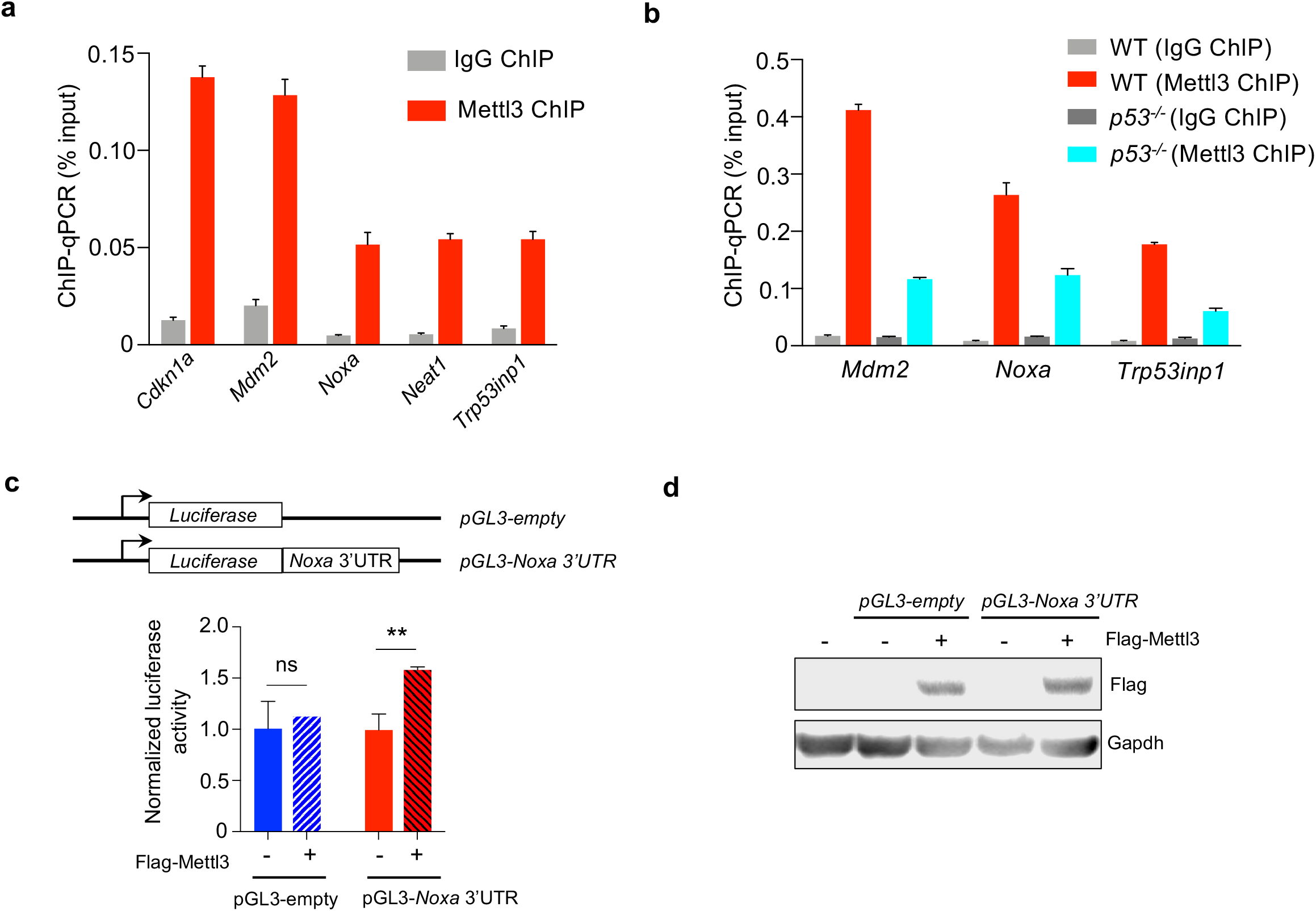
Mettl3 associates with chromatin of p53 target genes and enhances expression of *Noxa*. **a,** ChIP analysis of Mettl3 binding to p53 sites in p53 target gene loci, relative to input, in dox-treated *E1A;HRasV12* MEFs. IgG serves as a negative control antibody. Representative analysis from two biological replicates. **b,** ChIP analysis of Mettl3 binding to p53 sites in p53 target gene loci, relative to input, in dox-treated *E1A;HRasV12* wild-type and *p53^-/-^* MEFs. Representative analysis from two biological replicates. IgG serves as a negative control antibody, **c,** (Top) Reporter constructs expressing Firefly luciferase (Fluc) without and with the *Noxa* 3’UTR. (Bottom) Mean ± s.e.m of Fluc reporter activities in Flp-In 3T3 *Mettl3^-/-^* cells, transfected with empty vector or Flag-Mettl3 vector, after normalization to Renilla luciferase expression and subsequently to pGL3-empty without Mettl3 (n=3). *P* values were determined by unpaired, two-tailed Student’s *t*-test. \*\**P*<0.01, ns = not significant. **d.** Representative immunoblot showing Flag-Mettl3 protein levels in Flp-In 3T3 *Mettl3^-/-^* cells. Gapdh serves as a loading control (*n*=2).

### Mettl3 supports p53 in tumor suppression

In addition to serving as a key sentinel to genotoxic damage, p53 plays a critical role in responses to oncogenic signals, as underscored by its frequent mutation in human cancer^2^. In response to oncogene expression, p53 suppresses transformation *in vitro* and tumorigenesis *in vivo*. To determine whether Mettl3 might also contribute to p53 tumor suppressive function, we first utilized oncogene-expressing MEFs, a classical transformation model in which p53 potently suppresses transformation^24^. We knocked-down Mettl3 expression in oncogene-expressing MEFs using shRNAs. Analysis of these cells revealed that attenuated Mettl3 expression enhanced their clonogenic potential, a cardinal feature of transformed cells, supporting a role for Mettl3 in suppressing transformation (Fig. 6a). In contrast, Mettl3 knockdown in oncogene expressing *p53* null cells did not enhance clonogenic potential, indicating that the role for Mettl3 in transformation suppression is specifically in the context of intact p53 (Fig. 6b, Extended Data Fig. 5a). Interestingly, robust Mettl3 knockdown even appeared deleterious for viability in the absence of p53. We next assessed the consequences of Mettl3 knockdown for tumor growth by subcutaneous injection of oncogene-expressing cells into immunocompromised mice. Tumor growth in this context is significantly enhanced by p53 deficiency^25^, and, similarly, Mettl3 knockdown dramatically enhances tumor growth *in vivo*, suggesting a role for Mettl3 in tumor suppression (Fig. 6c). To more broadly assess the impact of Mettl3 deficiency on cancer development, we employed an autochthonous mouse lung adenocarcinoma model in which we could perform targeted gene inactivation using CRISPR/Cas9-mediated gene editing. In this model, lentiviral-Cre instillation induces oncogenic Kras, Cas9, and tdTomato expression as well as delivering sgRNAs targeting Mettl3 or a non-targeting control sgRNA. Importantly, p53 deficiency in this model is known to enhance tumor growth^24^. Interestingly, combined Mettl3 ablation by CRISPR/Cas9 and Kras activation resulted in enhanced lung adenocarcinoma growth relative to control mice (Fig. 6d, Extended Data Fig. 5b). Together, these findings suggest that Mettl3 has tumor suppressor activity in lung adenocarcinoma *in vivo*, in which p53 has an established role in suppressing cancer.

**Fig. 6.**
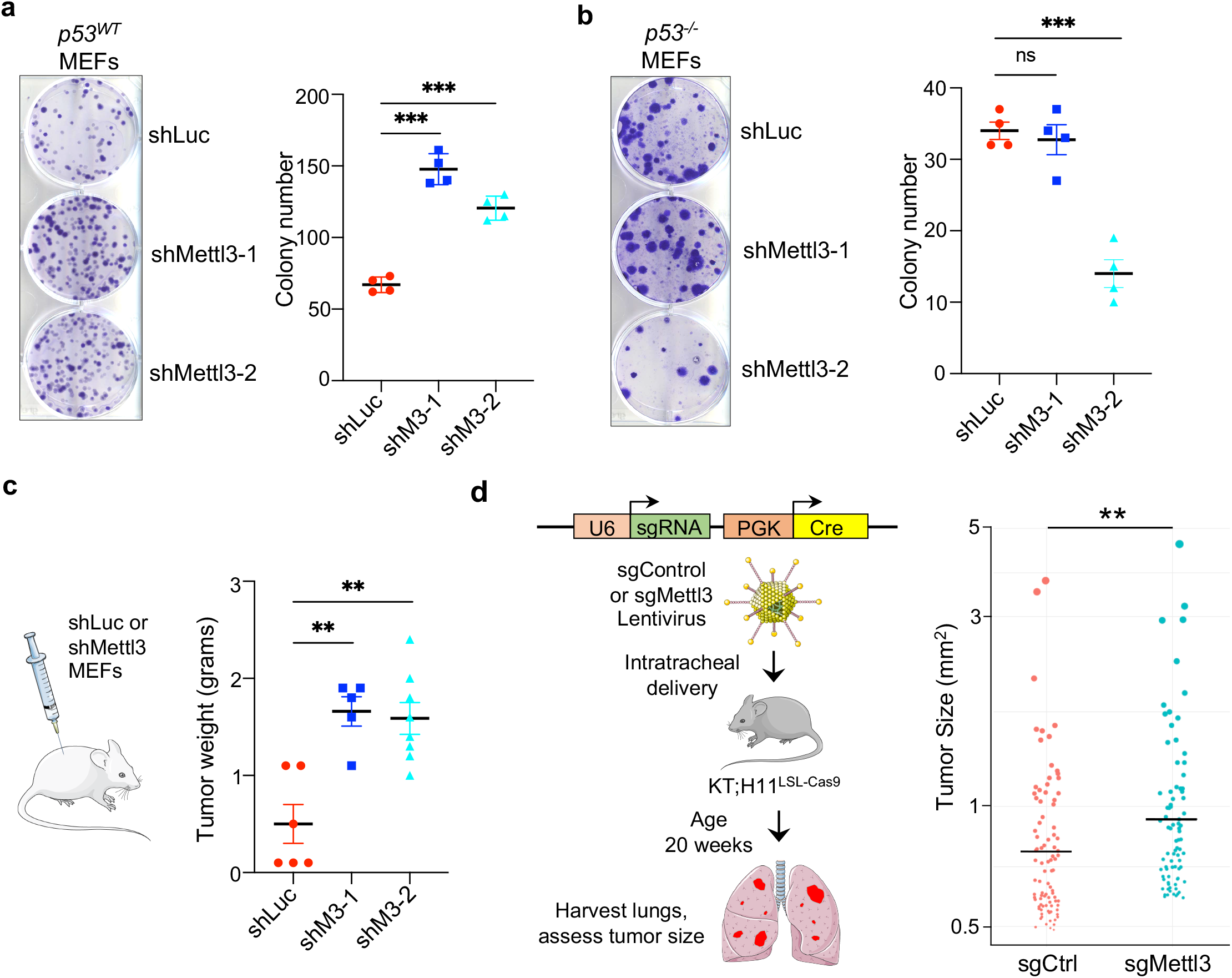
Mettl3 supports p53-mediated tumor suppression in mice. **a,b** Low density plating assay to assess clonogenic potential of *E1A;HRasV12*-expressing wild-type MEFs (**a**) or *p53^-/-^* MEFs (**b**) transduced with either shLuc or shMettl3 RNAs. (Left) Crystal violet was used to stain the colonies. Representative crystal-violet stained wells are shown. (Right) Average colony number (for *p53^WT^* MEFs, n=4, with triplicate samples, using two different MEF lines per genotype, for *p53^-/-^* MEFs, n=2 with triplicate samples). **c,** Average weight of *E1A;HRasV12* MEF tumors after growth *in vivo* for 32 days. Two different MEF lines were used. In **a,b**,**c** bar shows mean ± s.e.m. **d,** Lentiviral vectors expressing Cre recombinase and Mettl3 or control sgRNA were delivered intratracheally into *KT; H11^LSL- Cas9^* mice and tumor size of all lung tumors was assessed after 20 weeks (*n*=12 mice per group; *n*=907 control and 826 sgMettl3 tumors). Graph shows top 10% of all tumors in each group. Bar shows the mean for each group. *P* values were determined by unpaired, two-tailed Student’s *t*-test \*\**P*<0.01, \*\*\**P*<0.001, ns = not significant.

We next sought to interrogate the importance of a p53/methyltransferase axis in tumor suppression in human cancer. METTL3 is part of a multi-protein methyltransferase complex (MTC) that not only includes METTL14, but also WTAP, RBM15, RBM15b, VIRMA, CBLL1, and ZC3H13^26^. We thus sought to identify point mutations and deletions in *METTL3* complex components in human cancers and assess whether there is a mutually exclusive relationship with *TP53* mutations. Indeed, examination of the mutational spectrum in the full complement of MTC components in various carcinomas, including lung adenocarcinoma, breast, ovarian, and head and neck cancers, revealed a mutually exclusive pattern of mutations with *TP53*, suggesting that when constituents of the MTC are mutated, there is less selective pressure to mutate *TP53* (Figure 7a, Extended Data Fig. 6). We found further that like *TP53* mutation, MTC mutations compromise expression of various p53 target genes in a range of human cancer types, albeit not to the full extent seen with *TP53* mutation (Fig. 7b, Extended Data Fig. 7a). These data suggest that the MTC augments the p53 transcriptional program in humans. To test the functional significance of MTC activity in the p53 pathway, we leveraged DepMap data to assess the impact of *METTL3* knockout on proliferation of human cancer cell lines of different *TP53* status. Notably, we found that the most statistically significant *METTL3* co-dependency with all genes was with *TP53* (Extended Data Fig. 7b). Moreover, the highest Pearson correlation between Achilles scores with *METTL3* and *METTL14* knockout includes not only other MTC components, as expected, but also *TP53* and other positive regulators of the p53 pathway, *TP53BP1* and *ATM*, further supporting the idea that the MTC is a component of the p53 pathway (Fig. 7c, Extended Data Fig.7c). Conversely, both *METTL3* and *METTL14* showed a negative correlation with MDM2, as expected for a negative regulator of p53. Interestingly, the Achilles score for METTL3 is significantly lower in *TP53* mutant cell lines than in cell lines harboring wild-type *TP53* (Extended Data Fig. 7d), reminiscent of the reduced proliferation seen in oncogene-expressing, p53-deficient MEFs upon Mettl3 knockdown (Fig. 6b). Collectively, our findings support the idea that the METTL3 MTC and p53 operate in a common tumor suppressive pathway in various human cancers, with the MTC acting to promote full p53 activity.

**Fig. 7.**
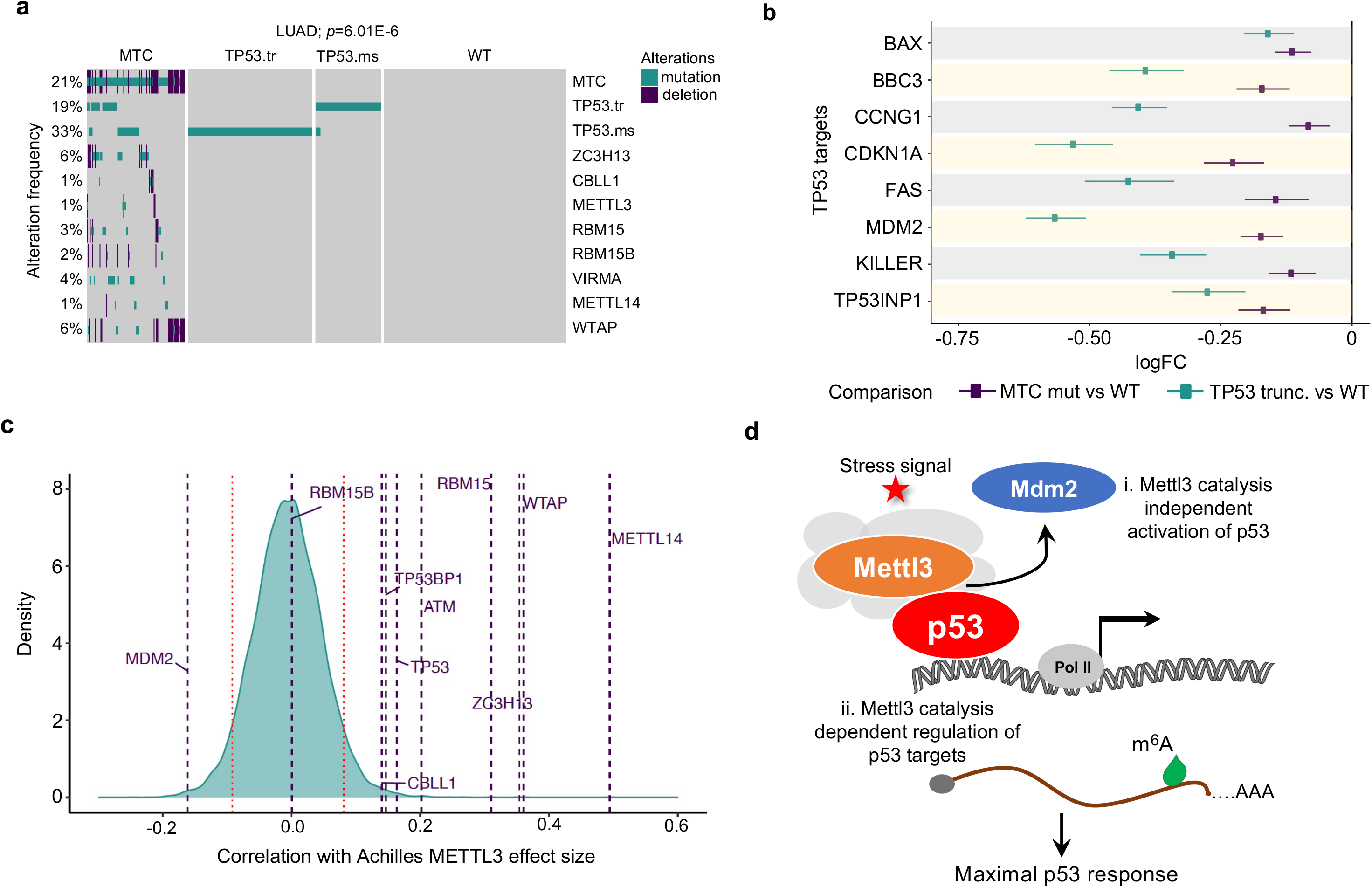
Mettl3 MTC and p53 operate in a common tumor suppressive pathway in human cancers. **a**, Oncoplot showing alteration frequencies of *TP53* and *METTL3* methyltransferase complex (MTC) components in human LUAD. TP53.tr refers to truncation mutations while TP53.ms refers to missense mutations in *TP53*. *P* value shows significance of DISCOVERY test unadjusted and adjusted for multiple testing. **b,** Differential expression (DE) analysis of *TP53* target genes in human cancers. Dots represent log fold change expression of select p53 targets in MTC mutant vs wild-type and *TP53* truncation mutant vs wild-type tumors. Summary represents DE in 33 TCGA cancer types. **c**, Density distribution of Pearson correlations between METTL3 Achilles scores. Horizontal lines represent genes of interest including MTC components, *TP53,* and regulators of p53 pathway. Red bars represent the 5^th^ and 95^th^ quantiles of the distribution. **d,** Proposed model of Mettl3-MTC regulation of the p53 pathway to potentiate full p53 responses to stress signals. In response to stress signals, Mettl3 stabilizes p53 protein in a m^6^A catalysis-independent manner by displacement of Mdm2 from p53 protein (i), and Mettl3 regulates the expression of select p53 pathway transcripts by governing their m^6^A-modification (ii).

## DISCUSSION

Here, we show that the Mettl3 complex plays a fundamental role in augmenting p53 activity at two levels: by binding to p53 to enhance its half-life through a catalytic activity-independent mechanism and by coordinating the m^6^A modification of p53 target gene transcripts in acute DNA damage responses and in tumor suppression to ensure their potent expression (Fig. 7d). This capacity ensures that a robust p53 response can be induced, and provides the potential for fine-tuning p53 activity.

Recruitment of the MTC by p53 and other sequence-specific transcription factors allows for co-transcriptional m^6^A modification that can provide an additional level of regulation for gene expression programs. A role for Mettl3 and the MTC in increasing activity of sequence-specific transcription factors has been reported, albeit through distinct mechanisms. For example, in AML, Mettl3 is recruited by the CEBPZ transcription factor to the promoters of oncogenes to drive m^6^A modification of emerging transcripts, a signal that ultimately augments translation of these transcripts to sustain AML cells^27^. In human pluripotent stem cells, in response to differentiation cues, the SMAD2/3 transcription factors recruit the MTC to RNAs to facilitate co-transcriptional m^6^A modification, leading to destabilization of specific transcripts such as *Nanog* to drive exit from pluripotency and promote differentiation^28^. Recruitment of the MTC to actively expressed genes in ES cells can also occur through Mettl14 binding to H3K36me3, a mark of transcription elongation, to drive co-transcriptional m^6^A modification of nascent mRNAs, also leading to destabilization of pluripotency gene mRNAs^29^. Thus, Mettl3 can be recruited to DNA by transcriptional regulators to promote m^6^A modification. Similarly, our findings suggest that Mettl3 is recruited to p53-bound sites in chromatin to co-transcriptionally install m^6^A marks on p53 target gene messages. Recruitment of the MTC by specific transcription factors may help to explain which transcripts are selected for m^6^A modification in a given setting, a point that has remained enigmatic.

m^6^A modification on RNA by writers like the MTC affects gene expression through a variety of mechanisms mediated by different m^6^A reader proteins^12^. While m^6^A marks can be read by YTHDF2/3 reader proteins to trigger mRNA destabilization, they can also be recognized by IGF2BP1-3 to promote mRNA stabilization^30–33^. m^6^A modification can also be recognized by the YTHDC1 nuclear reader to regulate splicing and RNA export^34^. In addition, m^6^A can affect translation through diverse mechanisms^12^. For example, Mettl3-driven m^6^A modification on the 5’UTR triggers binding to multiple subunits of the eukaryotic initiation factor 3 (eIF3) complex to promote cap-independent translation, while m^6^A marks in the CDS and 3’UTR results in YTHDF1/3 and eIF3 recruitment to promote translation initiation^32, 35, 36^. Interestingly, Mettl3 also promotes translation by acting as a reader: it can bind to an m^6^A mark in the 3’ UTR and interact with translation initiation machinery components such as eIF3H at the 5’ end of the transcript via RNA circularization, thus leading to enhanced translation of m^6^A modified mRNAs^37, 38^. In this capacity, Mettl3 catalytic activity is dispensable.

The Mettl3 complex is critical for responses to extracellular cues, either to promote cell fate transitions or homeostasis. For example, Mettl3 is required in mouse and human ES cells to restrict self-renewal and promote differentiation through downregulation of core pluripotency factor transcripts such as *Nanog*^16^. Similarly, Mettl3-mediated m^6^A modification in human hematopoietic stem cells is critical for differentiation *in vivo* and for inhibiting self-renewal in glioblastoma stem cells^39, 40^. Intriguingly, several studies have established a similar role of p53 in opposing stemness and promoting differentiation: p53 loss in mice triggers an expansion of both normal and cancer stem cells, and p53 restricts cellular reprogramming in iPS cell generation^41, 42^. Beyond modulating cell state, Mettl3 can also ensure homeostasis by promoting resolution to cellular stress. For example, in response to UV, Mettl3/Mettl14 localizes to DNA damage sites to transiently induce m^6^A RNA modification and promote DNA repair through recruitment of DNA polymerase^43^. Mettl3 also plays a critical role in the resolution of heat shock responses by inducing m^6^A modification on *Hsp70* mRNA, leading to its destabilization^44^. Our findings similarly underscore the importance of Mettl3 in promoting responses to signals, specifically for ensuring a robust p53 response to DNA damage or oncogene expression.

m^6^A modification and the MTC have a highly context-dependent role in cancer^45^. While Mettl3 has been reported to be an oncogene in numerous cancers such as AML, colon cancer, NSCLC, bladder cancer and ovarian cancer^27, 45, 46^, it has been shown to be a tumor suppressor in other settings, such as renal cell carcinoma and endometrial cancer^47, 48^. Similarly, Mett14 has been reported to promote some cancer types yet perform a tumor suppressive role in HCC and GBM^39, 45^. Our findings suggest that the MTC can be tumor suppressive in the context of intact p53, where it bolsters p53 activity. Interestingly, our clonogenic assays suggest that Mettl3 may support tumor growth in the absence of p53, as knockdown is detrimental for cell growth in this context. This difference in Mettl3 action in the context of active or deficient p53 provides one potential explanation for observed differences in the role of the MTC in cancer development. In addition, many studies of the MTC in cancer have relied on subcutaneous tumor xenograft studies in immunodeficient mice. It will be important to refine our understanding of the role of the MTC in cancer as performed here and in a previous AML study^49^ using autochthonous mouse models, which better mimic cancer initiation and progression in the context of an intact immune system and host stroma. Understanding the precise contexts in which Mettl3 must function to promote or suppress tumor development will be critical for better understanding pathways to tumorigenesis and for ultimately designing therapeutic interventions based on this pathway.

## FIGURE LEGENDS

**Extended Data Fig. 1| p53 interacts with Mettl3-Mettl14 methyltransferase complex independent of DNA. a,** Flp-In-3T3 *p53^-/-^* cells were co-transfected with Flag-Mettl3 and HA-p53, and the cell lysates were immunoprecipitated with anti-Flag M2 magnetic beads and detected by immunoblotting with the indicated antibodies (*n*=2). **b**, Co-IP and immunoblot assay to test interaction between endogenous p53 and Mettl14 in *E1A;HRasV12*-expressing MEFs. Flag antibody serves as non-specific antibody (*n*=2). **c,** Co-IP and immunoblot assay to test the nucleic acid dependence of interaction between endogenous p53 and Mettl3 in *E1A;HRasV12*-expressing MEFs. Lysates were pre-treated with ethidium bromide (EtBr) at 10 μg/ml prior to immunoprecipitations (*n*=2). **d,** Co-IP and immunoblot assay to test DNA-dependence of interaction between endogenous p53 and Mettl3 in lysates from MEFs pre-treated with DNaseI (40 U/ml) to degrade DNA. IgG serves as negative control antibody. IPs in *p53* null cells demonstrate the specificity of the p53 antibody (*n*=2). Representative immunoblots are shown in all panels.

**Extended Data Fig. 2| Mettl3 overexpression induces p53 protein levels and its deficiency negatively impacts p53 target gene induction under acute DNA damage. a,** Immunoblot after transfection of Flag-Mettl3 and HA-p53 plasmids into Flp-In-3T3 *p53^-/-^* cells. Gapdh serves as loading control (*n*=2). **b,** Immunoblots of wild-type and *Mettl3^-/-^* mouse ES cells showing expression of p53 and two of its canonical targets, p21 and Mdm2, in response to dox treatment. Gapdh serves as loading control (*n*=2). Representative immunoblots are shown in **a and b.**

**Extended Data Fig. 3| Mettl3 enhances p53 protein half-life under acute DNA damage.** Immunoblot analysis of p53 protein levels in mouse ES cells treated with doxorubicin (0.2 µg/ml for 6 hours) and cycloheximide (100 µM for indicated times). Gapdh serves as loading control. Data are representative of two biological replicates.

**Extended Data Fig. 4| Motif identification and distribution of m^6^A-peaks in *E1A;HRasV12*- expressing MEFs. a,** Top five sequence motifs enriched in m^6^A-modified mRNAs in MEFs expressing shLuc control or shMettl3 RNAs that were left untreated or treated with dox. **b,** Pie charts of the frequency distribution of m^6^A peaks that map to the listed mRNA features. m^6^A-IP reads were normalized to the total number of reads covering the m^6^A residue in the input.

**Extended Data Fig. 5.| Mettl3 supports p53 in colony formation and tumor suppression. a,** Immunoblotting for p53 and Mettl3 proteins in *E1A;HRasV12*-expressing *p53^-/-^* MEFs transduced with either shLuc or shMettl3 RNAs (*n*=1)**. b,** Representative images of H&E staining of lung tissue section from mice infected with Lenti-Cre sgControl (top panels) and sgMettl3 (bottom panels) viruses. Black scale bar = 500 μM, White scale bar = 50 μM.

**Extended Data Fig. 6| Mutual Exclusivity between *TP53* and METTL3 methyltransferase complex in human tumors. a,** Pan-cancer mutual exclusivity analysis using DISCOVER algorithm. MTC refers to any of the complex members while MTC (core) refers to METTL3, METTL14 and WTAP. **b,** Oncoplots showing alteration frequencies of METTL3-METTL14 methyltransferase complex components in human breast, ovarian and head & neck cancers. MTC refers to any of the complex members while MTC (core) refers to METTL3, METTL14 and WTAP. TP53.tr refers to truncation mutations while TP53.ms refers to missense mutations in *TP53*. *P* and *Q* values show significance of DISCOVERY test unadjusted and adjusted for multiple testing.

**Extended Data Fig. 7| *METTL3* and *TP53* operate in the same pathway. a,** Differential expression (DE) analysis of TP53 target genes in human cancers. Dots represent log fold change expression of select p53 targets in METTL3 complex (MTC) mutant vs wild-type and *TP53* truncation mutant vs wild-type tumors, in human lung adenocarcinoma (LUAD), breast cancer (BRCA), ovarian cancer (OV), uterine corpus endometrial carcinoma (UCEC) and head and neck squamous cell carcinoma (HNSCC). Summary represents DE in the five cancer types shown on the left. **b,** *METLL3* Achilles score association with all genes with any mutation across the Depmap (-log10 *P*-value). *P* values were calculated by a Wilcoxon signed-rank test. **c,** Density distribution of Pearson correlations between METTL14 Achilles scores. Horizontal lines represent genes of interest including MTC components, *TP53,* and regulators of p53 pathway. Red bars represent the 5^th^ and 95^th^ quantiles of the distribution. **d**, Achilles score for METTL3 is significantly lower (more essential, *P*=0.0031) in *TP53* mutant cell lines than in WT cell lines.

## METHODS

### Construction of Flp-In 3T3 p53-LAP, Flp-In 3T3 p53^-/-^ and Flp-In 3T3 Mettl3^-/-^ cell lines

We used Flp-In^TM^-3T3 cells (Thermo Fisher Scientific, Cat # R76107) to construct a cell line stably expressing C-terminally LAP-tagged wild-type p53 cDNA. Gateway entry vector for *Trp53* were created by BP recombination between pDONR221 and PCR amplified *Trp53* cDNA fragment. Flp-In system compatible C-terminally LAP-tagged p53 (p53-LAP) was generated by LR recombination between *Trp53* entry vector and pG-LAP7/puro destination vector (gift from Peter Jackson, Stanford University). Flp-In 3T3 cells stably expressing p53-LAP were generated by co-transfecting 0.4 µg of the preceding vector with 3.6 µg of pOG44, followed by selection with 4 µg/ml puromycin. Flp-In 3T3 p53^-/-^ cell line was generated by Crispr/Cas9 by co-transfecting Flp-In 3T3 cells with px330 p53 plasmid (Addgene Plasmid #59910) expressing Cas9 and sgRNA targeting mouse p53 and pmaxGFP plasmid (Lonza). Similarly, Flp-In 3T3 Mettl3^-/-^ cell line was generated using pX330 Mettl3 plasmid expressing Cas9 and sgRNA targeting mouse Mettl3. Two days post transfection, the GFP positive population was sorted by FACS and clonally expanded. Individual cell clones were screened for p53 or Mettl3 deletion using standard PCR and TIDE analysis. Loss of p53 or Mettl3 protein expression was confirmed by western blotting.

### Tandem Affinity Purification

After constructing Flp-In-3T3 expressing p53-LAP, we grew large scale cultures and affinity purified protein complexes for mass spectrometry. The cultures were either left untreated or treated with 0.2 µg/ml dox for 6 hours. Large-scale preparations of whole-cell lysates were subjected to dual-affinity purification, first with anti-GFP antibody-coupled beads to pull-down p53-LAP complexes. We then employed PreScission^TM^ protease, which cleaves at a unique site between the GFP and S-tags and performed a second round of affinity purification using a Protein S Agarose column that binds the S- tag. The bound p53-S-tag and any interacting proteins that were co-purified were eluted off the beads under denaturing conditions and run on a gradient gel, which was stained with Coomassie blue, and each lane was cut into 8 discrete bands, which were submitted for mass spectrometric protein identification. A 10 ml packed cell volume was re- suspended with 20 mL of LAP-resuspension buffer (300 mM KCl, 50 mM HEPES-KOH [pH 7.4], 1 mM EGTA, 1 mM MgCl2, 10% glycerol, 0.5 mM DTT, and protease inhibitors (Thermo Fisher Scientific, PI88266), lysed by gradually adding 0.6 mL 10% NP-40 to a final concentration of 0.3%, then incubated on ice for 10 min. The lysate was first centrifuged at 14,000 rpm (27,000 g) at 4°C for 30 min, and the resulting supernatant was centrifuged at 43,000 rpm (100,000 g) for 1 hr at 4°C to further clarify the lysate. High speed spin supernatant was mixed with 0.5 mL of GFP-coupled beads and rotated for 1 hr at 4°C to capture GFP-tagged proteins, and washed five times with 1 mL LAP200N buffer (200 mM KCl, 50 mM HEPES-KOH [pH 7.4], 1 mM EGTA, 1 mM MgCl2, 10% glycerol, 0.5 mM DTT, protease inhibitors, and 0.05% NP40). After re-suspending the beads with 1 mL LAP200N buffer lacking DTT and protease inhibitors, the GFP tag was cleaved by adding 5 mg of PreScission protease and rotating tubes at 4°C for 16 hours. All subsequent steps until the cutting of bands from protein gels were performed in a laminar flow hood to prevent keratin contamination. PreScission protease-eluted supernatant was added to 100 mL of S-protein agarose (EMD Millipore, 69704-3) to capture S-tagged protein. After washing three times with LAP200N buffer lacking DTT and twice with LAP100 buffer (100 mM KCl, 50 mM HEPES-KOH [pH 7.4], 1mM EGTA, 1mM MgCl2, and 10% glycerol), purified protein complexes were eluted with 50 µL of 2X LDS buffer and boiled at 95°C for 3 min. 5% of the total eluate was run on a gradient gel and silver-stained as quality control. Samples were then run on Bolt Bis-Tris Plus Gels (Thermo Fisher Scientific, NW04120BOX) in Bolt MES SDS Running Buffer (Thermo Fisher Scientific, B000202). Gels were fixed in 100 mL of fixing solution (50% methanol, 10% acetic acid in Optima LC/MS grade water (Thermo Fisher Scientific, W6-1) at room temperature, and stained with Colloidal Blue Staining Kit (Thermo Fisher Scientific, LC6025). After the buffer was replaced with Optima water, the bands were cut into eight pieces, followed by washing twice with 500 µL of 50% acetonitrile in Optima water. LC-MS/MS was performed by in-gel tryptic digestion of the gel bands followed by protein identification on a high performance Thermo Scientific Orbitrap Fusion™ Tribrid™ mass spectrometer as described below.

### Mass Spectrometry

Samples were processed for mass spectrometry by Stanford University Mass Spectrometry Facility. In a typical experiment, protein gel bands were first diced into 1 mm cubes and reduced with 5 mM DTT, 50 mM ammonium bicarbonate. After removal of residual solvent, proteins were alkylated using 10 mM acrylamide in 50 mM ammonium bicarbonate for 30 min at room temperature. Digestion was performed using Trypsin/LysC (Promega, Cat # V5071) in the presence of 0.02% Protease Max (Promega, Cat # V2071) overnight at 37°C. The following day, solid particulate was condensed by centrifugation and peptides extracted by adding 60% acetonitrile, 39.9% water, 0.1% formic acid and incubating for 15 min. Extracted peptides were dried in a speed vac and then reconstituted in 12.5 µl reconstitution buffer (2% acetonitrile with 0.1% Formic acid) and 3 µl of it was injected on the instrument.

Mass spectrometry experiments were performed using an Orbitrap Fusion™ Tribrid™ mass spectrometer (Thermo Scientific, San Jose, CA) with liquid chromatography performed using an Acquity M-Class UPLC (Waters Corporation, Milford, MA). For a typical LC MS experiment, a pulled-and-packed fused silica C18 reverse phase column was used, with Dr. Maisch 1.8-micron C18 beads as the packing material and a length of ∼25 cm. A flow rate of 450 nL/min was used with a mobile phase A of aqueous 0.2% formic acid and mobile phase B of 0.2% formic acid in acetonitrile. Peptides were directly injected onto the analytical column using a gradient (3-45% B, followed by a high-B wash) of 80 min. The mass spectrometer was operated in a data dependent fashion, with MS1 survey spectra collected in the orbitrap and MS2 fragmentation using CID for in the ion trap.

For data analysis, the .RAW data files were processed using Byonic (Protein Metrics, San Carlos, CA) to identify peptides and infer proteins. Proteolysis was assumed to be tryptic in nature and allowed for up to two missed cleavage sites. Precursor mass accuracies were held within 12 ppm, with MS/MS fragments held to a 0.4 Da mass accuracy. Proteins were held to a false discovery rate of 1%, using standard approaches^50^. Spectral counts from Byonic output were normalized by calculating NSAF values^51^ and bait - prey interactions were scored based on large number of unrelated affinity pull-downs in mouse cell lines^52^.

### Cell Culturing and drug treatments

Mouse embryonic fibroblasts, Flp-In-3T3 and H1299 cells were maintained in Dulbecco’s Modified Eagle Medium (Gibco) supplemented with 10% Fetal Calf Serum (FCS), 1% penicillin/streptomycin, 50 µg/mL gentamicin and incubated at 37°C in a carbon dioxide incubator. Mouse ES cells were cultured as previously described^16^. Doxorubicin (Sigma, Cat # D1515) treatment was done at a concentration of 0.2 µg/ml for 6 hours. Cycloheximide (Sigma, Cat # C7698) was treated at 100 µM for indicated period of time. Protein extracts were treated with DNase I (Invitrogen, Cat # 18047019) at 40 U/ml for 1 hr at 4°C. Ethidium Bromide was added at 10 µg/ml to protein extracts prior to and during IP and to the wash buffer.

### Plasmids

#### Cloning of p53 deletion mutants in pcDNA3.1 p53 backbone

Mutant p53 cDNAs were generated by polymerase chain reaction using pcDNA HA-p53 vector that contains N-terminal HA tagged full-length wild-type p53 as a template. To generate p53 (Δ1–42), the pair of primers used were: forward primer 5’- TTTTGGCGCGCCGATCTGTTGCTGCCCCAG-3’; and reverse primer: 5’-TTTTTTAATTAATCAGTCTGAGTCAGGCCC-3’. To generate p53 (Δ1–61), the pair of primers used were: forward primer 5’-TTTTGGCGCGCC CGAGTGTCAGGAGCTCCT- 3’; and reverse primer 5’-TTTTTTAATTAATCAGTCTGAGTCAGGCCC-3’.

#### Construction of pG-LAP2 Mettl3 (Flag-Mettl3)

Gateway entry vector for *Mettl3* were created by BP recombination between pDONR221 and PCR amplified *Mettl3* cDNA fragment. Flp-In system compatible N-terminally FLAG-tagged Mettl3 (Flag-Mettl3) was generated by LR recombination between *Mettl3* entry vector and pG-LAP2/puro destination vector (gift from Peter Jackson lab, Stanford University). Q5 site-directed mutagenesis (NEB E0554S) kit was used to generate the Flag-Mettl3 APPA mutant expression construct.

#### Construction of Lenti-U6-sgMettl3/Cre vector

We generated lentiviral vectors carrying PGK-Cre as well as an sgRNA targeting Mettl3 or control sgRNA targeting Neo. Lenti-U6-sgRNA/Cre vectors containing each sgRNA were generated as described previously ^53, 54^. Briefly, Q5 site-directed mutagenesis (NEB E0554S) kit was used to insert Mettl3 or control sgRNAs into the parental lentiviral vector containing the U6 promoter as well as PGK-Cre.

#### Construction of pX330 Mettl3 vector

We generated pX330 Mettl3 vector by cloning the previously described sgRNA targeting mouse Mettl3^16^ into the pX330 plasmid (Addgene Plasmid #42230). The pX330 plasmid was digested using BbsI and a pair of partially complementary annealed oligos containing overhangs from BbsI site and Mettl3 sgRNA sequence were cloned scarlessly into the vector. The oligo sequences used were: 5’-CACCGGGCTTAGGGCCGCTAGAGGT-3’ and 5’-AAACACCTCTAGCGGCCCTAAGCCC-3’.

#### Construction of pGL3-Noxa 3’UTR vector

A 311bp fragment encompassing the full exon 3 and a portion of the 3’UTR of mouse *Noxa* gene that harbors a Mettl3-dependent m^6^A motif was cloned into the XbaI site immediately downstream of the *luciferase* gene in pGL3 vector using conventional cloning strategy. The primers used for PCR amplification of the Noxa 3’UTR fragment were: forward primer 5’-AATCTAGAGACTTGAAGGACGAGTGT-3’; and reverse primer 5’- AATCTAGATTCACGTTATCACAGCTC-3’.

### Co-immunoprecipitation assays

Cells were harvested by scraping method using cell scrapers (Corning, Cat # 3010) and lysed with ice cold NP-40 lysis buffer (50 mM Tris pH 8.0, 150 mM NaCl, 1% NP-40, 0.5 mM EDTA, 10% Glycerol) containing protease inhibitors (Roche cOmplete^TM^, Cat # 11 697 498 001). Protein was quantitated using the bicinchoninic acid protein assay (BCA) kit (Pierce, Cat # 23227). 1-2 mg total protein was used for each immunoprecipitation reaction (IP) reaction in 500 µl final volume. Lysates were first pre-cleared using 50% slurry of BSA blocked Protein A sepharose beads (GE, Cat # 17-0780-01) by incubating for 30 minutes at 4°C. For Mettl3 and Mettl14 IPs, the pre-cleared lysates were incubated with 1-2 µg Mettl3 polyclonal antibody (Abclonal, A8370) or 1-2 µg Mettl14 polyclonal antibody (Abcam, ab98166) overnight at 4°C on a nutator to allow protein complexes to form. For p53 IPs, pre-cleared lysates were incubated with 2-4 µl p53 polyclonal antibody (Leica Biosystems, NCL-L-p53-CM5p) overnight at 4°C on a nutator to allow p53 protein complexes to form. The day after, immune complexes were retrieved with 50 µl of 50% slurry of BSA blocked Protein A sepharose beads for 4 hours at 4°C. Post the incubation, the beads were washed 3 times using 0.1% NP-40 containing wash buffer (50 mM Tris pH 8.0, 150 mM NaCl, 0.1% NP-40, 0.5 mM EDTA, 10% Glycerol). The immobilized immunoprecipitated complexes were eluted by boiling the sepharose beads in 2X SDS sample buffer. For Flag-HA CoIPs, lysates were pre-cleared using Protein A/G magnetic beads (Thermo Fisher Scientific, Cat # 26162) for 30 minutes at 4°C . The pre-cleared lysates were incubated with 25 µl Flag M2 magnetic beads (Millipore, Cat # M8823) for 4 hours at 4°C to immunoprecipitate Flag-tagged Mettl3. Beads were washed 4 times using 1% Triton X-100 containing wash buffer (10 mM Tris pH 8.0, 150 mM NaCl, 1% Triton X- 100, 1 mM EDTA, 1 mM EGTA). Flag-protein complexes were eluted using Flag peptide (Millipore, Cat # F3290) at 150 µg/ml by incubating the beads at room temperature for 30 minutes. The eluates were resolved on a 10% SDS-PAGE gel and the proteins were electroblotted onto PVDF membranes (Millipore, Immobilon-P, Cat # IPVH20200) for probing with following primary and secondary antibodies: anti-p53 (gift from Helin K, Univ. of Copenhagen, clone AI25, 1:500), anti-Mettl3 (Abclonal, A8370, 1:500), anti-Mettl14 (Abcam, ab98166), anti-Mdm2 (Abcam, ab16895, 1:500), anti-Flag (Sigma, F1804, 1:1000) and anti-HA (Thermo Fisher Scientific, 71-5500, 1:500), peroxidase Affinipure goat anti-mouse IgG, light chain specific (Jackson ImmunoResearch, 115-035-174, 1:5,000), peroxidase IgG fraction monoclonal mouse anti-rabbit IgG, light chain specific (Jackson ImmunoResearch, 211-032-171, 1:5,000). Inputs represent 2-5% of the lysate subjected to immunoprecipitation.

### Immunoblotting

Protein was extracted using NP-40 lysis buffer (50 mM Tris, pH 8.0, 150 mM NaCl, 1% NP-40, 0.5 mM EDTA, and 10% glycerol) containing protease inhibitors (Roche cOmplete^TM^, Cat # 11 697 498 001). Protein was quantitated using the BCA kit (Pierce, Cat # 23227). 20 μg of protein was resolved on a 10% SDS-PAGE gel, electroblotted onto PVDF membranes (Millipore, Immobilon-P, Cat # IPVH20200) and blocked in 5% non-fat dry milk prepared in TBS with 0.1% Tween-20 (TBST). Three washes were performed in TBST, and the following primary and secondary antibodies were used: rabbit anti-p53 (Leica Biosystems, NCL-L-p53-CM5p, 1:5000), rabbit anti-Mettl3 (Abclonal, A8370, 1:1000), rabbit anti-Mettl14 (Abcam, ab98166, 1:1000), rabbit anti-p21 (Abcam, ab188224, 1:1,000), mouse anti-Mdm2 (Abcam, ab16895, 1:1000), mouse anti-Flag (Sigma, F1804, 1:1000) and rabbit anti-HA (Thermo Fisher Scientific, 71-5500, 1:500), mouse anti-Gapdh (Fitzgerald, 10R-G109A, 1:10,000), peroxidase Affinipure goat anti-rabbit IgG (H+L) (Jackson ImmunoResearch, 111-035-144, 1:5,000), or peroxidase Affinipure goat anti-mouse IgG (H+L) (Jackson ImmunoResearch, 115-035-003, 1:5,000). Immunodetection was performed using ECL^TM^ Prime (Millipore-Sigma, Cat# GERPN2232) or Clarity^TM^ Western ECL substrate (Bio-Rad, Cat# 1705060).

### qRT-PCR

RNA extraction was performed using Trizol reagent (Thermo Fisher Scientific, Cat

#15596018) according to the manufacturer’s protocol. RNA (2-5 µg) was treated with DNAse I (Thermo Fisher Scientific, Cat # AM1906) according to the manufacturer’s instructions. Reverse transcription was conducted with M-MLV reverse transcriptase (Thermo Fisher Scientific, Cat # 28025) and random primers (Thermo Fisher Scientific, Cat # 48190). 1 µg of total RNA was used for cDNA synthesis. cDNA was diluted 1:5 in nuclease-free water and stored at –80°C until used. Quantitative PCR was performed in triplicate using PowerUP SYBR green master mix (Thermo Fisher Scientific, Cat # A25743) and a 7900HT Fast Real-Time PCR machine (Applied Biosystems). Expression analysis was performed using specific primers for each gene (Extended Data Table 1). The mean of housekeeping gene *β-Actin* was used as an internal control to normalize the variability in expression levels. All qRT-PCR performed using PowerUP SYBR Green was conducted at 50°C for 2 min, 95°C for 10 min, and then 40 cycles of 95°C for 15 s and 60°C for 1 min. Melt curve analysis was done to verify the specificity of the reaction. Samples were quantified using a standard curve.

**Extended Data Table 1.**
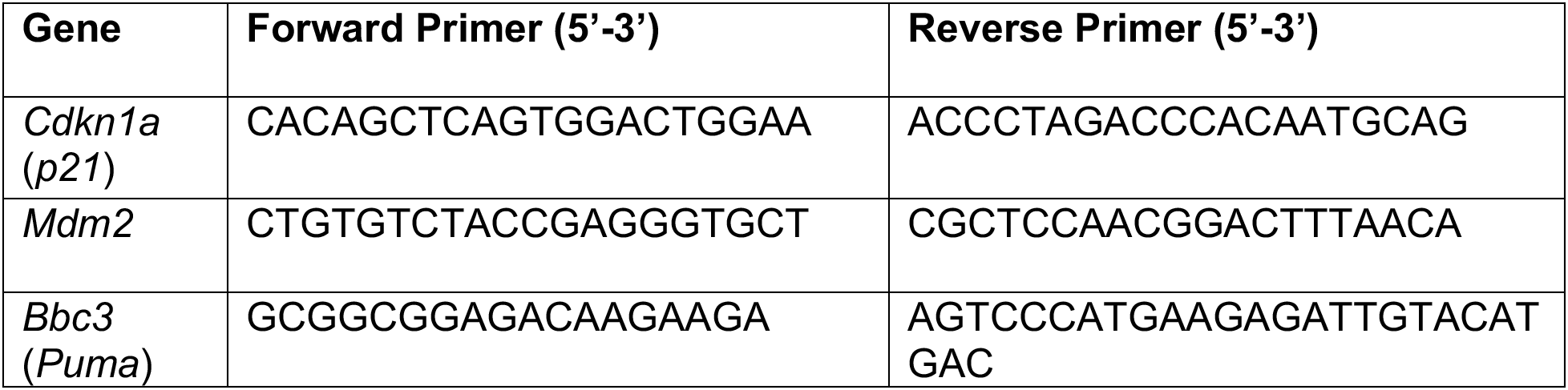
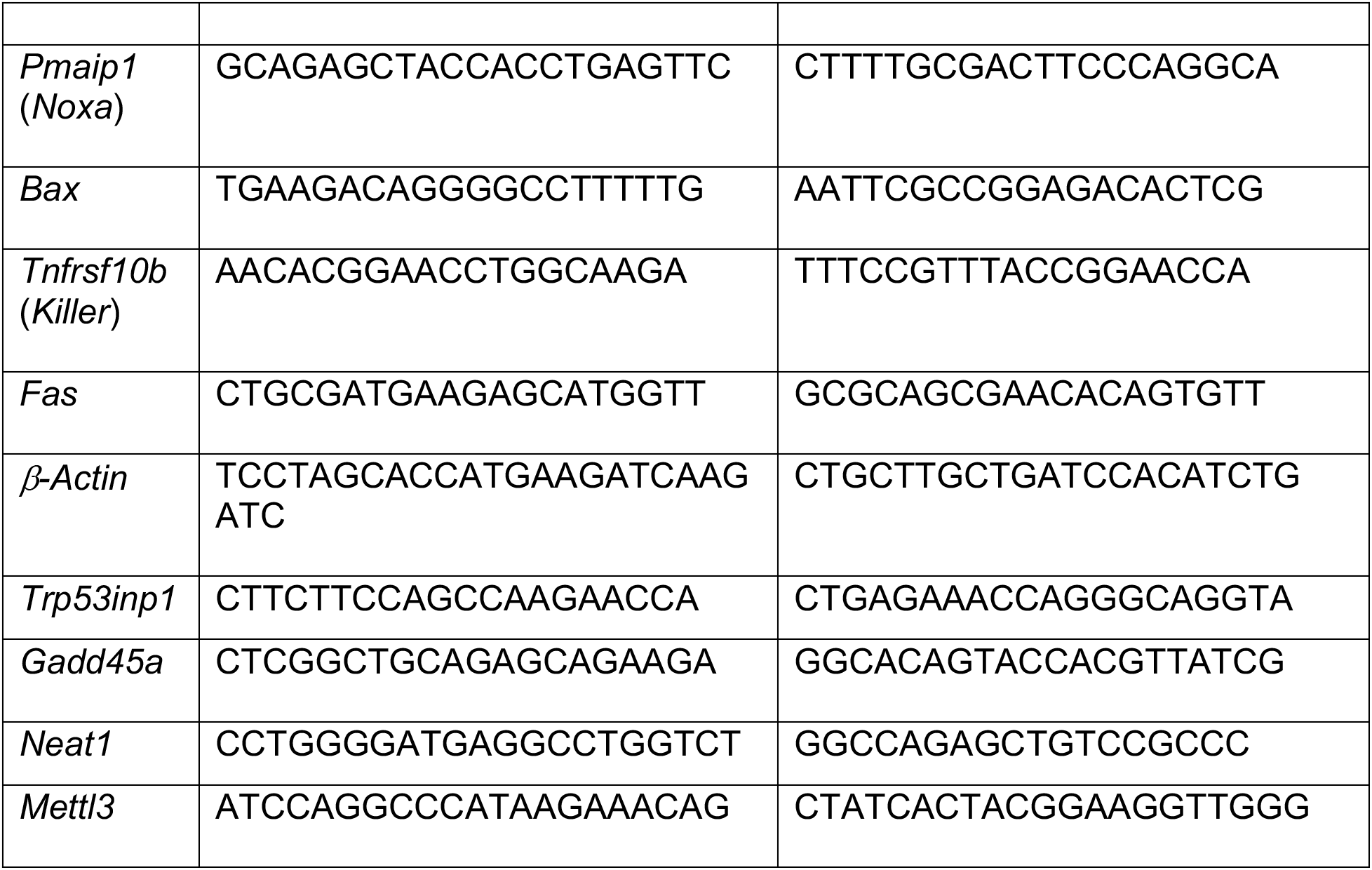
qRT-PCR primer list

### ChIP

Analysis of Mettl3 chromatin binding was done in Wild-type and *p53^-/-^ E1A;HRasV12*- expressing MEFs. MEFs were seeded at 7 x 10^6^ cells per 10 cm dish, one day prior to the ChIP experiment. After treatment with 0.2 μg/ml doxorubicin for 6h, cells were harvested to prepare chromatin for immunoprecipitation using either p53 polyclonal antibodies (Leica Biosystems, Cat # NCL-L-p53-CM5p) or Mettl3 polyclonal antibodies (Abclonal, Cat # A8853). ChIPs were performed essentially as described previously (Kenzelmann Broz, D. et al, 2013)^23^. Chromatin-immunoprecipitated DNA was analyzed by quantitative PCR with binding site-specific primers (Extended Data Table 2) using PowerUP SYBR green master mix (Thermo Fisher Scientific, Cat # A25743) and a 7900HT Fast Real-Time PCR machine (Applied Biosystems). The signals obtained from the ChIP were analyzed by the percent input method.

**Extended Data Table 2.**
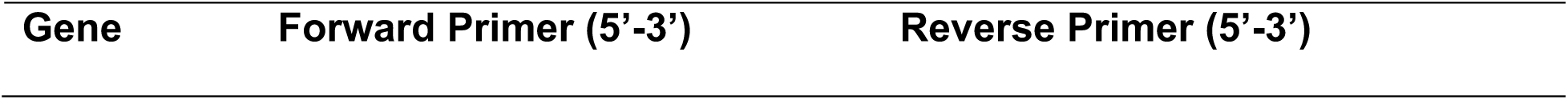
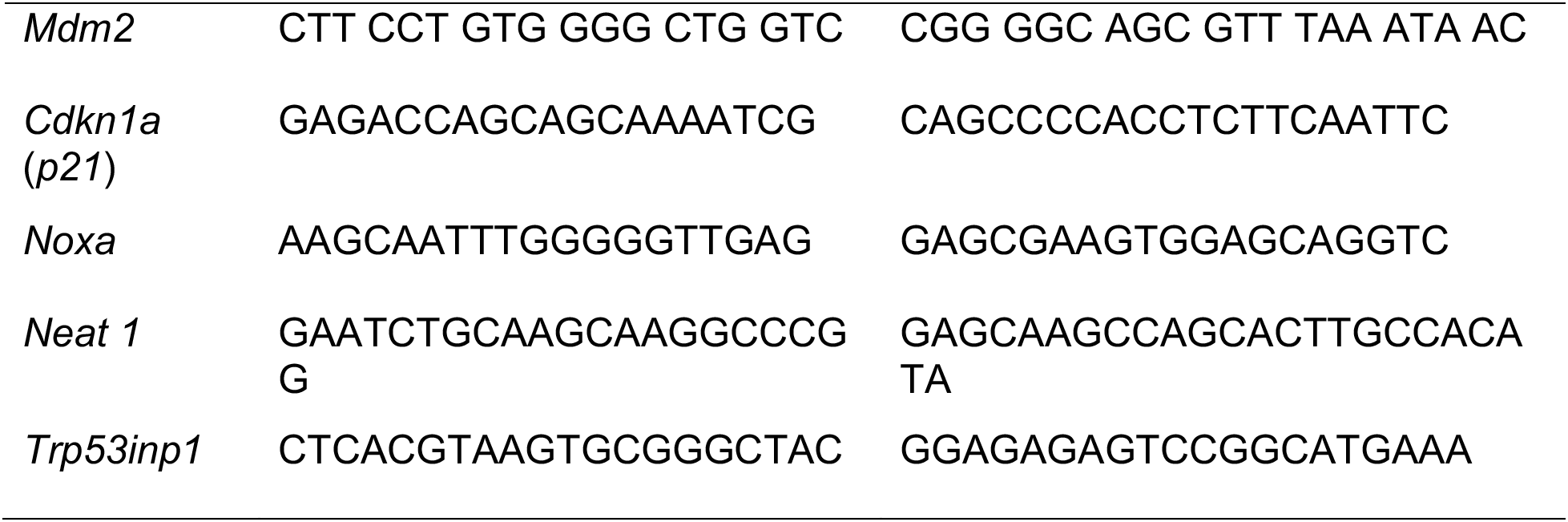
ChIP-qPCR primer list

### Lentiviral shRNA cell lines

pSico shMettl3 contructs were a gift from Pedro Batista, NIH. Lentivirus carrying shMettl3 were produced by co-transfecting 293T cells with 150 ng of pCMV-VSV-G, 350 ng of pCMV-dR8.2 dvpr, and 500 ng of pSico shMettl3 plasmids previously described. Media was replaced 16 hr after transfection to omit transfection reagent, and virus was harvested at 48 hr post-transfection. Virus was then filtered with a 0.45 µm PVDF filter (SLHV013SL, Millipore) and mixed with polybrene (TR-1003-G, Millipore). Wild-type or p53 null *E1A;HRasV12*-expressing MEFs^24^ were infected for 48 hr, followed by selection with 2 µg/ml puromycin.

### m^6^A-eCLIP-seq

m^6^A-eCLIP was performed at Eclipse BioInnovations Inc, San Diego CA as described below. The procedure started by isolating mRNAs through poly(A) selection using oligo (dT) beads (Thermo Fisher Scientific) followed by magnetic bead separation from 50 μg of total RNA. The mRNA was DNase treated at 37°C for 10 minutes then immediately sheared into 100-200nt fragments by heating at 95°C for 12 minutes in 1x Turbo DNase buffer (Thermo Fisher Scientific). An anti-m^6^A antibody (CST) was added, and samples were UV-C crosslinked using a UVP CL-1000 crosslinker at 2 rounds of 150 mJ/cm2 using 254nm wavelength. The antibody-RNA complexes were then coupled overnight to protein G beads (CST). Following overnight coupling, library preparation (including adapter ligations, SDS-PAGE electrophoresis and nitrocellulose membrane transfer, reverse transcription, and PCR amplification) was performed as previously described for standard eCLIP^55^, with the 30-110 kDa region size-selected by cutting from the membrane (corresponding to RNA fragments crosslinked to antibody heavy and light chains). 10 ng of fragmented mRNA was run as an RNA-seq input control, starting with FastAP treatment as described^55^. The final library’s shape and yield was assessed by Agilent TapeStation.

The eCLIP cDNA adapter contains a sequence of 10 random nucleotides at the 5′ end. This random sequence serves as a unique molecular identifier (UMI)^56^ after sequencing primers are ligated to the 3′ end of cDNA molecules. Therefore, eCLIP reads begin with the UMI and, in the first step of analysis, UMIs were pruned from read sequences using umi_tools (v0.5.1)^57^. UMI sequences were saved by incorporating them into the read names in the FASTQ files to be utilized in subsequent analysis steps. Next, 3′-adapters were trimmed from reads using cutadapt (v2.7) [https://doi.org/10.14806/ej.17.1.200] and reads shorter than 18 bp in length were removed. Reads were then mapped to a database of mouse repetitive elements and rRNA sequences compiled from Dfam^58^ and Genbank^59^. All non-repeat mapped reads were mapped to the mouse genome (mm10) using STAR (v2.6.0c)^60^. PCR duplicates were removed using umi_tools (v0.5.1) by utilizing UMI sequences from the read names and mapping positions. Peaks were identified within m^6^A-eCLIP samples using the peak caller CLIPper (https://github.com/YeoLab/clipper)61. For each peak, IP versus input fold enrichment values were calculated as a ratio of counts of reads overlapping the peak region in the IP and the input samples (read counts in each sample were normalized against the total number of reads in the sample after PCR duplicate removal). A p-value was calculated for each peak by the Yates’ Chi-Square test, or Fisher Exact Test if the observed or expected read number was below 5. Comparison of different sample conditions was evaluated in the same manner as IP versus input enrichment; for each peak called in IP libraries of one sample type we calculated enrichment and p-values relative to normalized counts of reads overlapping these peaks in another sample type. Peaks were annotated using transcript information from GENCODE (Release M21)^62^ with the following priority hierarchy to define the final annotation of overlapping features: protein coding transcript (CDS, UTRs, intron), followed by non-coding transcripts (exon, intron). The RPM value for a particular region was calculated by counting the number of reads in that region and dividing that by the “per million” scaling factor, which is defined as the total number of reads in the sample divided by 1,000,000. The log2 fold changes for a peak are calculated as follows: Log2 (RPM in IP in peak region / RPM in input in peak region). Log2 fold change values (RPM in IP in peak region / RPM in input in peak region) were calculated for individual peaks in each m^6^A-IP to normalize the IP reads to the total number of reads covering the m^6^A residue. The differential of Log2 fold change between shLuc and shMettl3 samples treated with dox, was used to identify genes that were most enriched for a specific m^6^A modification under DNA damage in a Mettl3- dependent manner.

### Luciferase reporter assays

pGL3-empty or pGL3-Noxa 3’UTR *luciferase* constructs were co-transfected with pRL- SV40 (*Renilla* plasmid) and pgLAP2-*Mettl3* into Flp-In 3T3 *Mettl3^-/-^* cells. Luciferase activity was measured 24 hours after transfection. Values were first normalized to internal control, Renilla luciferase and then to control sample (pGL3-empty vector with no Mettl3 reconstitution). Immunoblots were performed in parallel samples to confirm the expression of Flag-Mettl3.

### Low plating assays

shLuc and shMettl3 *E1A;HRas^V12^* MEFs were plated on six-well plates at 150 cells per well in triplicate and grown for 10-12 days. Cells were fixed with 10% formalin and stained with 0.1% crystal violet stain. Plates were scanned, and colonies were counted manually.

### Mice

ICR SCID male mice were obtained from Taconic Biosciences (Cat # ICRSC-M). *Kras^LSL- G12D^* (K), *Rosa26^LSL-Tomato^* (T), and *H11^LSL-Cas9^* (Cas9) mice have been described previously^53, 63, 64^. All mice were maintained under pathogen-free conditions at the Stanford animal care facility. All experiments were approved by Administrative Panel on Laboratory Animal Care at Stanford University.

### Subcutaneous tumor assays

Subcutaneous tumor studies were performed as described previously^25^. Briefly, 1 × 10^6^ shLuc or shMettl3 *E1A;Ras^V12^* MEFs were subcutaneously injected into the flanks of Scid mice, and tumors were weighted at 32 days post-injection.

### Lung Adenocarcinoma assay

Lentiviral particles were produced and tittered as described previously^54^. Lung tumors were initiated by intratracheal infection of mice as described previously^65^ using lentiviral- Cre vectors at the indicated titers. Lung tumors were induced by intratracheal administration of 90,000 particles of Lenti-U6-sgRNA/PGK-Cre virus to 6-12 week old mice as previously described^65^. Lungs were harvested at 20 weeks after inoculation and tumor burden was assessed by histology as indicated.

### Histology

Hematoxylin and eosin (H&E) staining on paraffin embedded lung tissues were performed using standard protocols. A NanoZoomer 2.0-RS slide scanner (Hamamatsu) was used for imaging.

### Mutual Exclusivity, Differential Expression Analysis and Achilles Perturbation Data

TCGA gene expression data was downloaded from gdc.cancer.gov. Whole Exome Sequencing annotations (MAF file) was downloaded from^66^. Gene expression raw counts were normalized using *voom* from *limma* package. Limma was used to calculate the TP53 targets differential expression between TP53 truncated tumors and TP53 wild-type, and between MTC altered (mutated, amplified or deleted) tumors and MTC wild-type for each tumor type individually. Summary fold changes were obtained using the function *combine.est* from *genefu* package.

Discover test^67^ was used to calculate mutual exclusivity across MTC genes (mutated or deleted) and TP53 (missense or truncated mutations) across all TCGA cancer tissues individually and combined (adjusted by tissue).

Achilles perturbation scores and CCLE mutations were downloaded from DepMap site (depmap.org, version 20Q2). Two sample Wilcoxon test was used to compare METTL3 Achilles score and mutation profiles (hotspot mutations as defined in DepMap). The Mettl3 Achilles scores were scaled such that the mean is 0 and the standard deviation is 1. Scaled scored were calculated using the formula: scaled score = (actual score-mean (actual score))/sd(actual score). Pearson correlation was used to calculate correlation across Achilles scores.

## ACKNOWLEDGEMENTS

We thank Pedro Batista for the kind gift of the pSico shMettl3 constructs and James Broughton for bioinformatics support. This work was supported by an NIH R35 grant (CA197591) and the Virginia and D.K. Ludwig Fund for Cancer Research to LDA. H.Y.C. is an Investigator of the Howard Hughes Medical Institute.

## AUTHOR CONTRIBUTIONS

L.D.A. conceived and designed the overall study. N.R, M.W, N.A.M, J.D., A.M.K., J.A.S., T.B., A.S.M, C.M. performed and analyzed experiments. All authors contributed to the interpretation of experiments. L.D.A and N.R. wrote and edited the manuscript. All authors reviewed the manuscript.

