## Supplemental Figures for "The Mettl3 epitranscriptomic writer amplifies p53 stress responses"

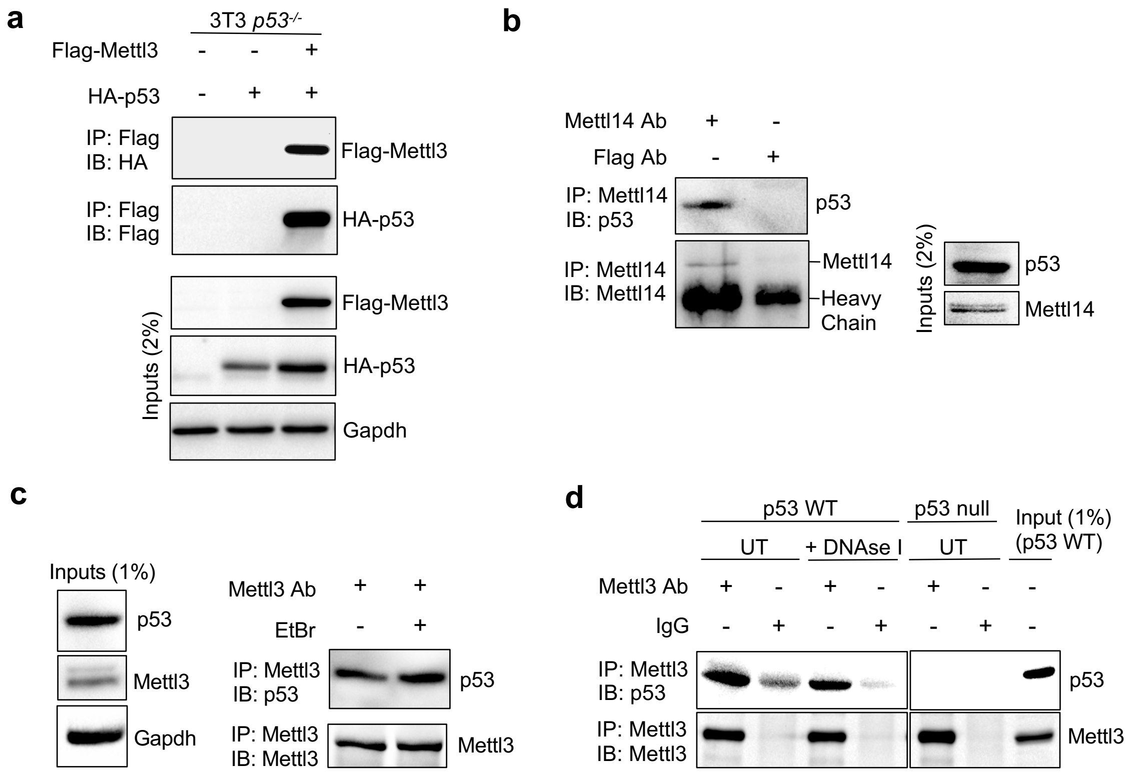


**Extended Data Fig. 1| p53 interacts with Mettl3-Mettl14 methyltransferase complex independent of DNA. a,** Flp-In-3T3 *p53*-/- cells were co-transfected with Flag-Mettl3 and HA-p53, and the cell lysates were immunoprecipitated with anti-Flag M2 magnetic beads and detected by immunoblotting with the indicated antibodies (*n*=2). **b**, Co-IP and immunoblot assay to test interaction between endogenous p53 and Mettl14 in *E1A;HRasV12*-expressing MEFs. Flag antibody serves as non-specific antibody (*n*=2). **c,** Co-IP and immunoblot assay to test the nucleic acid dependence of interaction between endogenous p53 and Mettl3 in *E1A;HRasV12*-expressing MEFs. Lysates were pre-treated with ethidium bromide (EtBr) at 10 μg/ml prior to immunoprecipitations (*n*=2). **d,** Co-IP and immunoblot assay to test DNA-dependence of interaction between endogenous p53 and Mettl3 in lysates from MEFs pre-treated with DNaseI (40 U/ml) to degrade DNA. IgG serves as negative control antibody. IPs in *p53* null cells demonstrate the specificity of the p53 antibody (*n*=2). Representative immunoblots are shown in all panels.

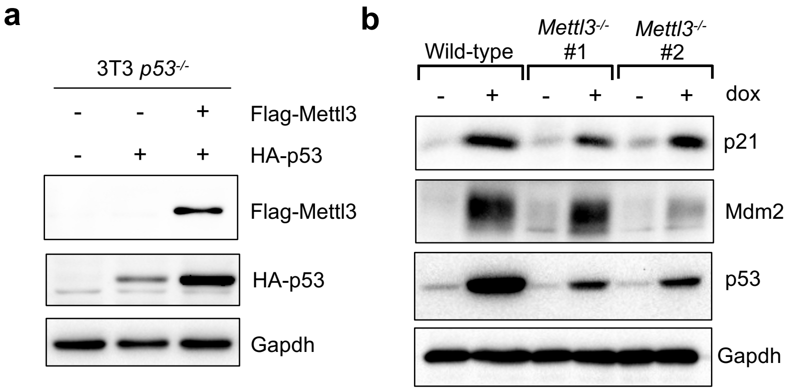


**Extended Data Fig. 2| Mettl3 overexpression induces p53 protein levels and its deficiency negatively impacts p53 target gene induction under acute DNA damage. a,** Immunoblot after transfection of Flag-Mettl3 and HA-p53 plasmids into Flp-In-3T3 *p53*$^{-/-}$ cells. Gapdh serves as loading control (*n*=2). **b,** Immunoblots of wild-type and *Mettl3*$^{-/-}$ mouse ES cells showing expression of p53 and two of its canonical targets, p21 and Mdm2, in response to dox treatment. Gapdh serves as loading control (*n*=2). Representative immunoblots are shown in **a** and **b.**

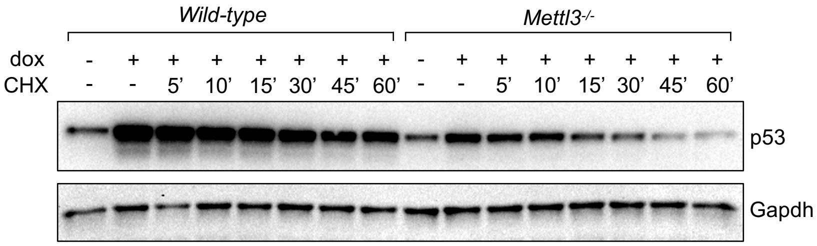


**Extended Data Fig. 3| Mettl3 enhances p53 protein half-life under acute DNA damage.** Immunoblot analysis of p53 protein levels in mouse ES cells treated with doxorubicin (0.2 µg/ml for 6 hours) and cycloheximide (100 µM for indicated times). Gapdh serves as loading control. Data are representative of two biological replicates.

**a**

shLuc (UT)

| Rank | Motif | *P*-value | % of Targets |
|---|---|---|---|
| 1 | 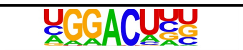 | 1E-1163 | 62.16% |
| 2 | 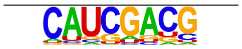 | 1E-236 | 23.99% |
| 3 | 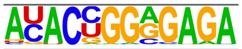 | 1E-202 | 0.79% |
| 4 | 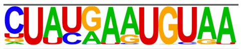 | 1E-193 | 0.60% |
| 5 | 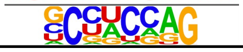 | 1E-134 | 28.01% |

shMettl3-2 (UT)

| Rank | Motif | *P*-value | % of Targets |
|---|---|---|---|
| 1 | 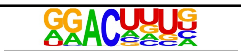 | 1E-1083 | 70.38% |
| 2 | 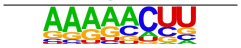 | 1E-248 | 24.29% |
| 3 | 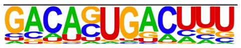 | 1E-245 | 20.13% |
| 4 | 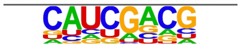 | 1E-232 | 45.67% |
| 5 | 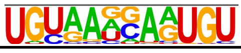 | 1E-174 | 1.69% |

shLuc (dox)

| Rank | Motif | *P*-value | % of Targets |
|---|---|---|---|
| 1 | 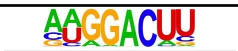 | 1E-916 | 71.6% |
| 2 | 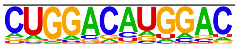 | 1E-287 | 29.89% |
| 3 | 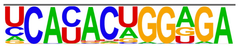 | 1E-280 | 1.59% |
| 4 | 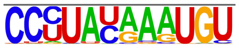 | 1E-278 | 1.66% |
| 5 | 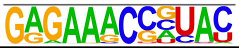 | 1E-263 | 2.08% |

shMettl3-2 (dox)

| Rank | Motif | *P*-value | % of Targets |
|---|---|---|---|
| 1 | 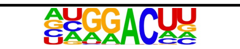 | 1E-1110 | 65.75% |
| 2 | 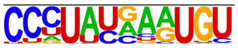 | 1E-320 | 1.00% |
| 3 | 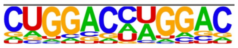 | 1E-279 | 38.57% |
| 4 | 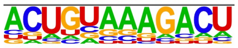 | 1E-223 | 28.63% |
| 5 | 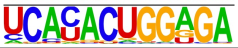 | 1E-189 | 1.25% |

**b**

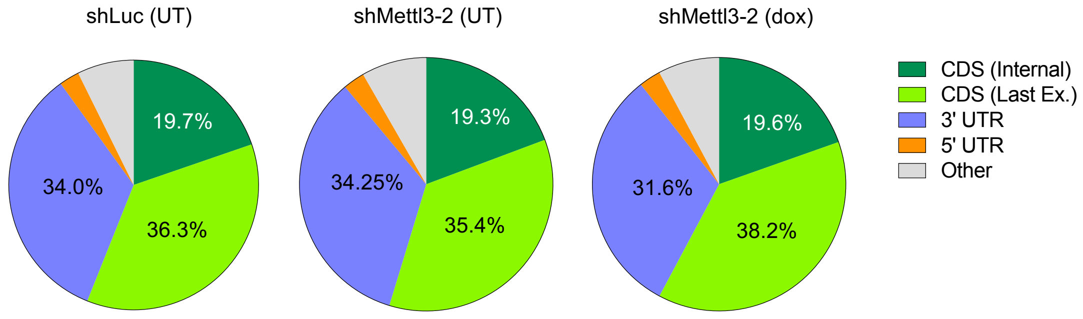


**Extended Data Fig. 4| Motif identification and distribution of m⁶A-peaks in *E1A;HRasV12*- expressing MEFs. a,** Top five sequence motifs enriched in m⁶A-modified mRNAs in MEFs expressing shLuc control or shMettl3 RNAs that were left untreated or treated with dox. **b,** Pie charts of the frequency distribution of m⁶A peaks that map to the listed mRNA features. m⁶A-IP reads were normalized to the total number of reads covering the m⁶A residue in the input.

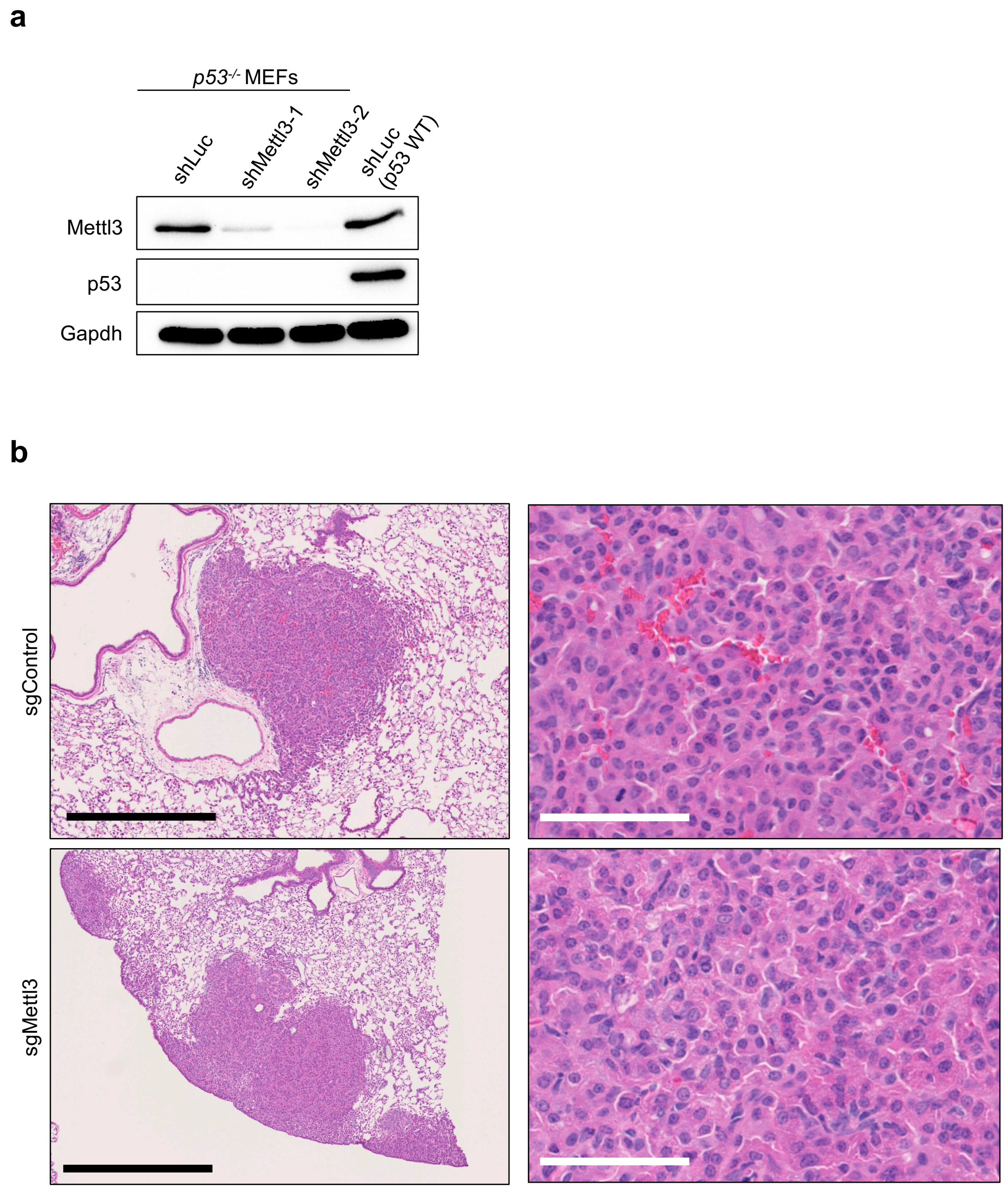


**Extended Data Fig. 5.| Mettl3 supports p53 in colony formation and tumor suppression. a,** Immunoblotting for p53 and Mettl3 proteins in *E1A;HRasV12*-expressing *p53*$^{-/-}$ MEFs transduced with either shLuc or shMettl3 RNAs (*n*=1)**. b,** Representative images of H&E staining of lung tissue section from mice infected with Lenti-Cre sgControl (top panels) and sgMettl3 (bottom panels) viruses. Black scale bar = 500 μM, White scale bar = 50 μM.

**a**

Pan-cancer ME

| gene | *P* value | *Q* value |
|---|---|---|
| MTC | 4.99E-14 | 5.48E-13 |
| ZC3H13 | 1.35E-11 | 6.93E-11 |
| RBM15 | 3.57E-07 | 1.35E-06 |
| MTC (core) | 2.19E-06 | 7.4E-06 |
| RBM15B | 9.79E-06 | 2.72E-05 |
| VIRMA | 2.26E-05 | 5.83E-05 |
| METTL14 | 0.000145 | 0.000326 |
| WTAP | 0.000631 | 0.001361 |
| METTL3 | 0.001886 | 0.003998 |
| CBLL1 | 0.00275 | 0.00535 |

**b**

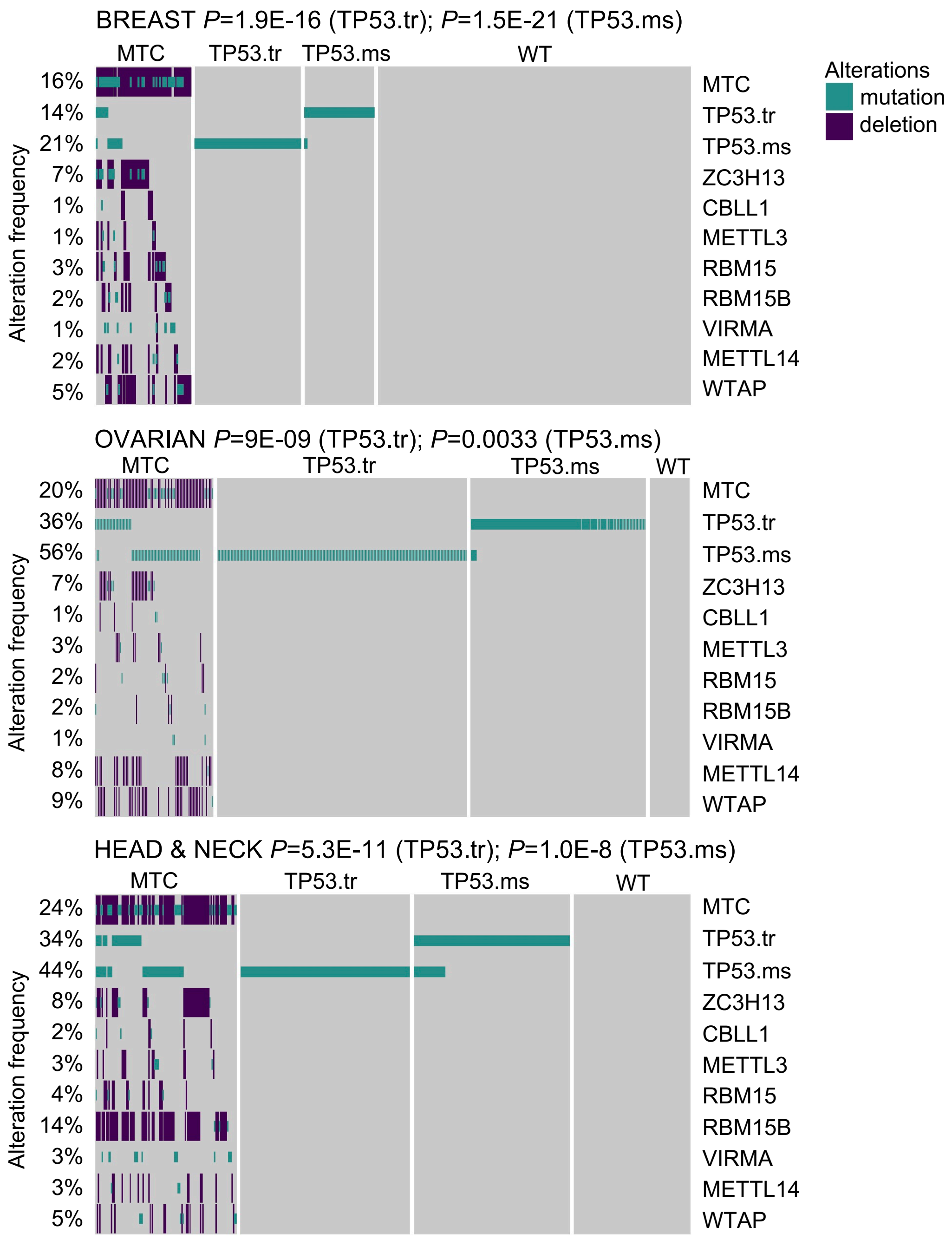


**Extended Data Fig. 6| Mutual Exclusivity between *TP53* and METTL3 methyltransferase complex in human tumors. a,** Pan-cancer mutual exclusivity analysis using DISCOVER algorithm. MTC refers to any of the complex members while MTC (core) refers to METTL3, METTL14 and WTAP. **b,** Oncoplots showing alteration frequencies of METTL3-METTL14 methyltransferase complex components in human breast, ovarian and head & neck cancers. MTC refers to any of the complex members while MTC (core) refers to METTL3, METTL14 and WTAP. TP53.tr refers to truncation mutations while TP53.ms refers to missense mutations in *TP53*. *P* and *Q* values show significance of DISCOVERY test unadjusted and adjusted for multiple testing.

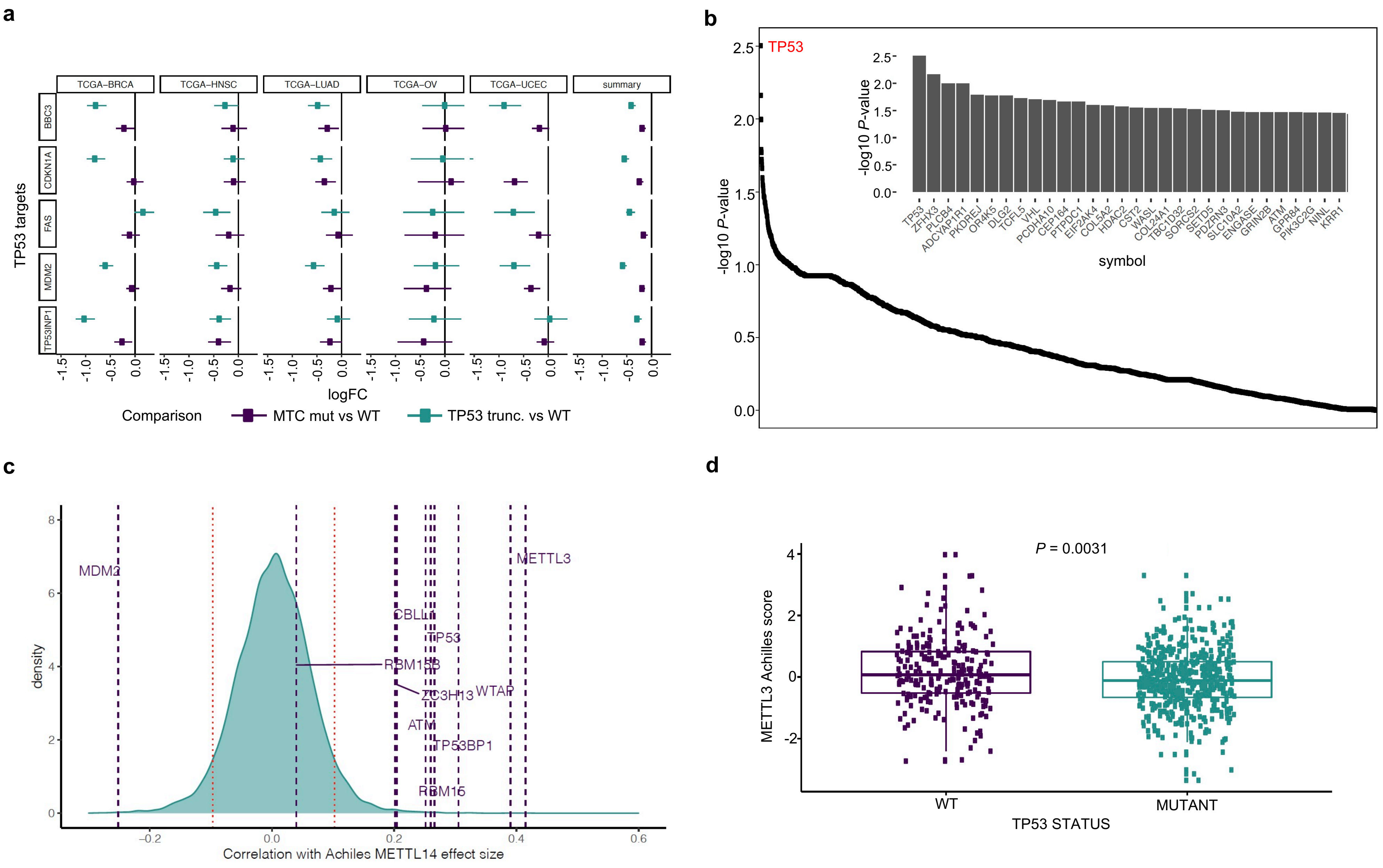


**Extended Data Fig. 7| *METTL3* and *TP53* operate in the same pathway. a,** Differential expression (DE) analysis of TP53 target genes in human cancers. Dots represent log fold change expression of select p53 targets in METTL3 complex (MTC) mutant vs wild-type and *TP53* truncation mutant vs wild-type tumors, in human lung adenocarcinoma (LUAD), breast cancer (BRCA), ovarian cancer (OV), uterine corpus endometrial carcinoma (UCEC) and head and neck squamous cell carcinoma (HNSCC). Summary represents DE in the five cancer types shown on the left. **b,** *METLL3* Achilles score association with all genes with any mutation across the Depmap (-log10 *P*-value). *P* values were calculated by a Wilcoxon signed-rank test. **c,** Density distribution of Pearson correlations between METTL14 Achilles scores. Horizontal lines represent genes of interest including MTC components, *TP53,* and regulators of p53 pathway. Red bars represent the $5^{th}$ and $95^{th}$ quantiles of the distribution. **d,** Achilles score for METTL3 is significantly lower (more essential, *P*=0.0031) in *TP53* mutant cell lines than in WT cell lines.
